# Enhancing adoptive T cell therapy with synergistic host immune engagement promotes long-term protection against solid tumors

**DOI:** 10.1101/2022.09.10.507437

**Authors:** Kwasi Adu-Berchie, Joshua M. Brockman, Yutong Liu, David K.Y. Zhang, Alexander J. Najibi, Alexander Stafford, Miguel C. Sobral, Yoav Binenbaum, Maxence O. Dellacherie, David J. Mooney

## Abstract

Adoptive T cell therapy provides the T cell pool needed for immediate tumor debulking, but the infused T cells generally have a narrow repertoire for antigen recognition and limited ability for long-term protection. Here, we present a biomaterial platform that enhances adoptive T cell therapy by synergistically engaging the host immune system via in-situ antigen-free vaccination. T cells alone loaded into these localized cell depots provided significantly better control of subcutaneous B16-F10 tumors than T cells delivered through direct peritumoral injection or intravenous infusion. The anti-tumor response was significantly enhanced when T cell delivery was combined with biomaterial-driven accumulation and activation of host immune cells, as this prolonged the activation state of the delivered T cells, minimized host T cell exhaustion, and enabled long-term tumor control. This integrated approach provides both immediate tumor debulking and long-term protection against solid tumors, including against tumor antigen escape.

## Introduction

Adoptive T cell therapy is an emerging area of immunotherapy that involves the ex-vivo manipulation of T cells and their subsequent reinfusion into patients^1^. The T cells are either genetically modified autologous polyclonal T cells isolated from apheresed PBMCs, or tumor infiltrating lymphocytes (TILs), which are harvested from tumors, expanded ex vivo and reinfused into patients^2^. Adoptive T cell therapy has shown remarkable promise in hematological cancers, but has seen limited efficacy in solid tumors^3^. Inspired by reports that localized, intra-tumoral injection of T cells yield superior therapeutic outcomes^4,5^, recent efforts have focused on developing biomaterial-based T cell depots which can be implanted or injected at the tumor site to further enhance therapeutic T cell efficacy^6–10^. While adoptively transferred T cells provide the T cell source needed for immediate tumor debulking, their long-term efficacy can be hindered by their narrow repertoire of antigen recognition, which can result in tumor antigen escape, T cell exhaustion and limited long-term protection^11–13^.

We hypothesized that a system which simultaneously enhances adoptive T cell therapy and effectively engages host immunity can provide immediate tumor debulking and long-term protection against solid tumors. To address this hypothesis, we developed a “Synergistic In-situ Vaccination Enhanced T cell depot” (SIVET) to serve as an integrated platform for locally delivering adoptively transferred T cells to the tumor site, while accumulating and activating large numbers of host antigen presenting cells (APCs) to process and present antigens from the dying tumor cells to generate host anti-tumor T cell responses (Fig 1a). First, the therapeutic efficacy of the T cells alone loaded into the localized depot was compared with systemic delivery and direct peritumoral T cell delivery, with or without a sub-lethal irradiation pre-conditioning regimen. Next, the ability of two SIVET formulations, concentrating host APCs via release of either FLT3L or GM-CSF, to elicit host T cell responses was characterized, as well as the crosstalk between the adoptively transferred T cells and host immune cells. Finally, the therapeutic efficacy of SIVET was investigated, including in a model of tumor antigen escape.

**Fig. 1.**
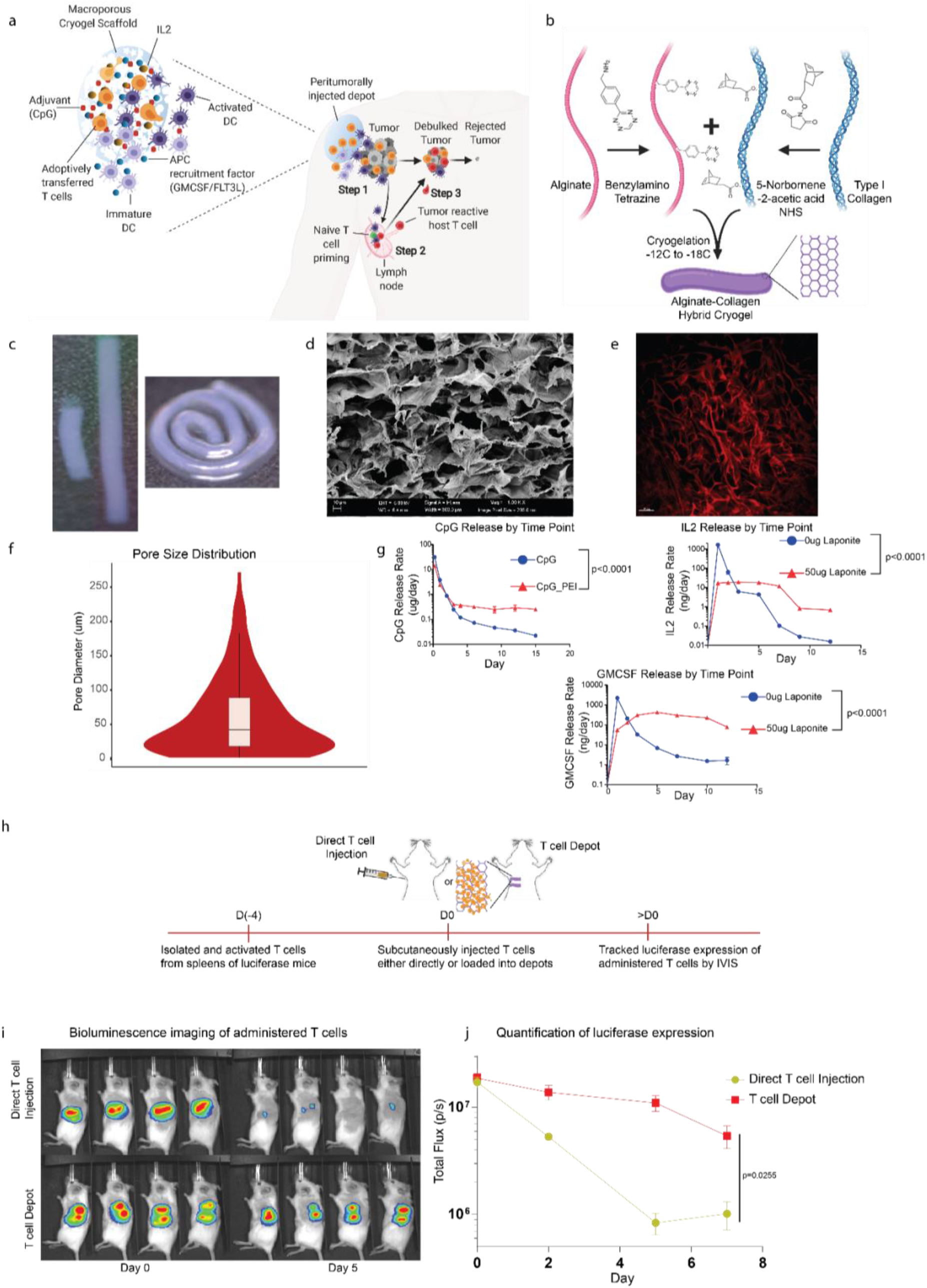
Synthesizing and characterizing SIVETs. a. Schematic detailing the proposed mechanism of action. **Step 1:** Depots infused with T cells and loaded with factors that recruit antigen presenting cells (APC) are injected adjacent to a tumor. Depots concurrently release T cells to debulk the tumor while recruiting APCs to the tumor site, where they become activated and process tumor antigen. **Step 2:** Tumor-antigen presenting APCs migrate to lymph nodes and prime host naïve, antigen-specific T cells. **Step 3:** Primed tumor reactive host T cells traffic to the tumor, which is undergoing tumor debulking by the adoptively transferred T cells, and facilitate tumor rejection and long-term anti-tumor immunity. b. Schematic of SIVET fabrication. Alginate and collagen type 1 are modified with tetrazine and norbornene respectively, and then reacted under cryogelation conditions (−12C to −18C) to form macroporous alginate-collagen hybrid cryogels. c. Photographs of rod-shaped cryogels showing tunable lengths (left) and flexibility (right). d. SEM image of cryogel showing macroporous structure. e. SHG demonstrating pristine cryogel architecture and pore-size distribution. f. Quantification of pore-size distribution from SHG represented as a violin plot. g. Tunable release of immunomodulatory factors from cryogels: CpG with or without PEI condensation (top left), IL2 and GM-CSF with or without pre-adsorption onto laponite (top right and bottom respectively). p-values for g-i were determined by two-way ANOVA with repeated measures. Data are mean ± s.e.m from n=3. h-j. Analysis of local T cell persistence in vivo. h. Schematic of experimental set-up. CD8+ T cells were isolated from spleens of luciferase expressing mice, activated in vitro, and either directly injected subcutaneously or loaded into depots before injection. i. Bioluminescence images of administered T cells over time. j. Quantification of T cell luciferase expression over time. p-value was determined by two-way ANOVA with repeated measures. Data are mean ± s.e.m from n=4.

## Results

### Synthesizing and characterizing SIVETs

Synergistic In-situ Vaccination Enhanced T cell depot (SIVETs) were fabricated by modifying alginate and type I collagen with tetrazine and norbornene respectively, which react bio-orthogonally via inverse electron demand Diels–Alder click reaction^14^. Under cryogelation conditions, this reaction yields macroporous cryogels with shape recovery properties that readily enable needle injection (Fig 1b). The collagen provides ligands for T cell adhesion and migration^15^ while the alginate provides structural support to the depot. The depots are rod-shaped, which minimized shear during injection and enhanced scalability, as the length of the depots could be easily tuned (Fig 1c). The depots are also highly flexible, allowing them to conform to the area of injection (Fig 1c). Scanning electron microscopy and second harmonic imaging confirmed that the depots contained pores with an average size of 50um, with collagen comprising the pore walls (Fig 1d-f). Changing the ratio of alginate to collagen significantly altered the ability of the depots to recover their shape after deformation, while the total percentage of polymer influenced their interconnected porosity (Supplementary Fig 1a-d).

To assess the ability of SIVETs to release soluble immunomodulatory factors, the cytokines IL2 and GMCSF, which facilitate T cell expansion and myeloid cell recruitment respectively^16,17^, and the adjuvant CpG, which activates myeloid cells^18^, were loaded into depots by adding the factors to the gel mixture before cryogelation. The interactions of these bioactive factors with the depot was altered either by adsorption onto highly negatively charged laponite nanoparticles (for cytokines)^19^ or by polyethyleneimine condensation (for CpG)^20^ prior to loading (Supplementary Fig 1e), resulting in tunable release profiles for all three factors (Fig 1g). Immunomodulatory factors were loaded into depots in concert with laponite or PEI for subsequent studies.

To establish the need for adhesion motifs, T cells were subsequently loaded into either alginate-only cryogels or alginate-collagen hybrid depots and their migration analyzed. T cells loaded into alginate-collagen hybrid depots showed significantly faster migration speed and longer track lengths than T cells loaded into alginate-only cryogels (Supplementary Fig 2).

### T cell depots enhance local T cell persistence in vivo and provide superior tumor control

Local in vivo T cell persistence following administration of cell loaded depots was next investigated. CD8+ T cells from the spleens of luciferase expressing mice^21,22^ were activated and either directly injected subcutaneously into mice with soluble IL2 (Direct T cell Injection) or infused into T cell depots with matched IL2 quantity before injection (T cell Depot). IVIS images and subsequent quantification showed that while similar amounts of T cells were delivered on Day 0, there were significantly more luciferase expressing CD8+ T cells at the site of injection at subsequent timepoints with T cell depots (Fig 1h-j).

The therapeutic efficacy of cells administered via T cell depots was subsequently examined in the aggressive B16-F10 tumor model after preconditioning by sub-lethal irradiation, a common clinical procedure to improve T cell engraftment in adoptive therapies^23^. CD8+ T cells isolated from pmel mice, which recognize the gp100 antigen on B16-F10 tumor cells^24,25^, were administered to tumor bearing mice 5 days after B16-F10 inoculation (Fig 2a). For preconditioning, mice were subjected to total body sub-lethal irradiation on day 4 after tumor inoculation. Tumor growth and subsequent mouse survival showed that peritumoral injection of T cell depots infused with pmel CD8+ T cells resulted in significant enhancement of tumor control over intravenously (IV) injected T cells and direct peritumoral T cell injection (Fig 2b-c). Importantly, the peritumoral injection of empty depots did not affect tumor growth.

**Fig. 2.**
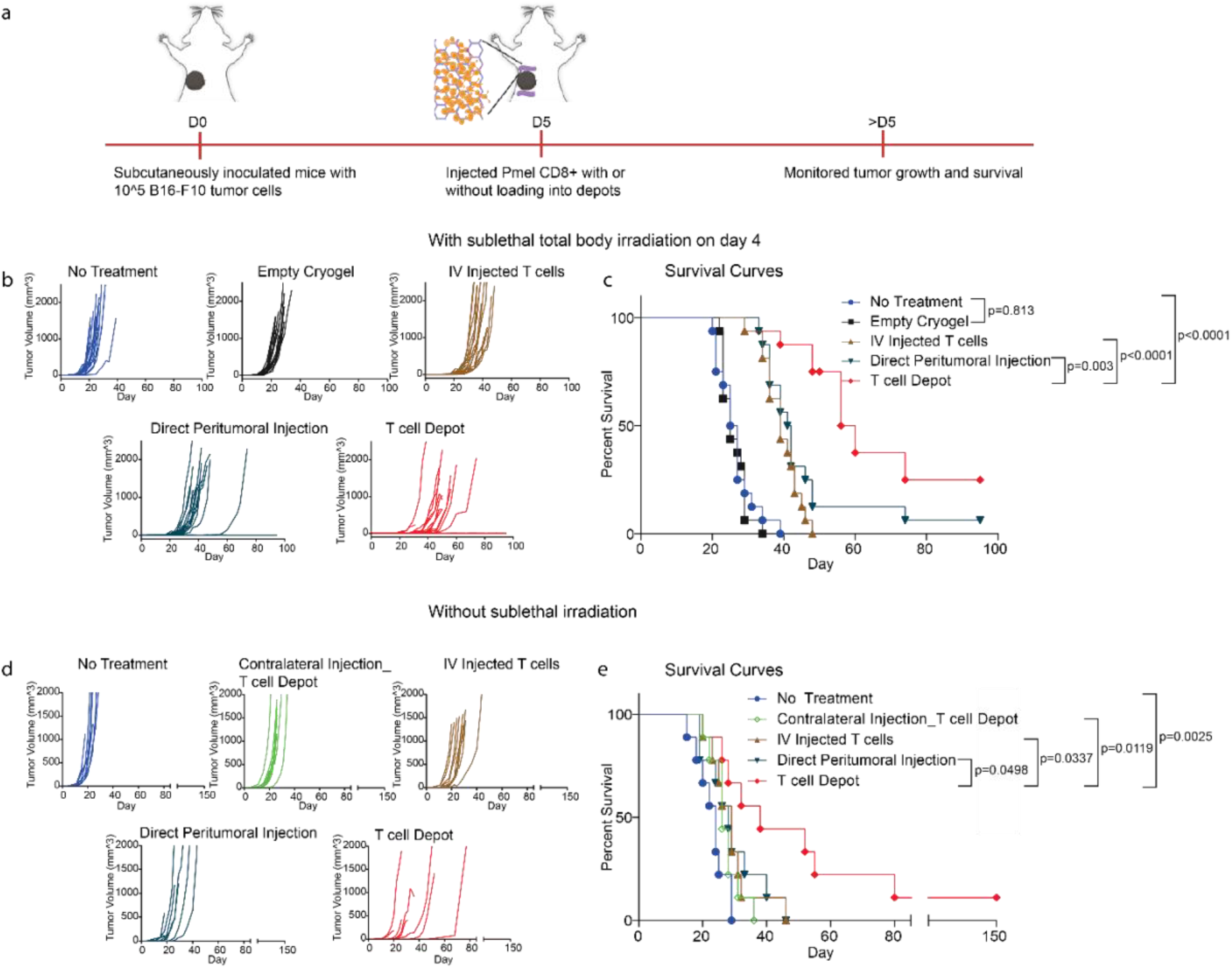
T cell depots alone enhance B16-F10 tumor control. a. Schematic of therapeutic study. Tumor volumes (b) and Kaplan-Meier survival curves (c), with sub-lethal irradiation preconditioning of mice left untreated, injected peritumorally with empty depots or treated with the following T cell conditions: intravenous T cell delivery, direct peritumoral T cell injection or T cell depots. p-values for c were determined by Log-rank (Mantel-Cox) test. Data are n=15-16 mice per condition, 2 independent studies (7-8 mice per study). d-e. Therapeutic study without preconditioning. Tumor growth (d) and Kaplan-Meier survival curves (e) of the indicated conditions. For the ‘Contralateral Injection_T cell Depot’ condition, T cell loaded depots were injected contralateral to the tumor location. p-values for e were determined by Log-rank (Mantel-Cox) test. Data are n=7 or 8 mice per condition.

Next, the ability of T cell-only depots to control B16-F10 tumors without sub-lethal irradiation was investigated, to assess the ability of T cell depots to enable transferred T cell engraftment without prior preconditioning. T cells administered by IV or direct peritumoral injection showed significantly diminished tumor control when pre-conditioning was not performed. However, T cell-only depots still showed superior control of B16-F10 tumors (Fig 2d-e). The therapeutic effect was significantly reduced when the T cell depots were injected contralateral to the tumor site (Fig 2d-e), confirming the significance of localized T cell delivery. Subsequent studies were performed without sublethal irradiation.

### SIVETs elicit host T cell responses

SIVET both locally delivers T cells and concentrates and activates host antigen presenting cells (APCs) to the tumor site for antigen uptake and both local antigen presentation and migration to lymph nodes for host T cell priming. After establishing that T cell depots by themselves resulted in better therapeutic outcomes compared with traditional modes of T cell delivery, we sought to understand the impact of the complete SIVET on both adoptively transferred T cells and host immune cells. The relative fractions of tumor reactive host T cells were first analyzed in lymph nodes and spleens 9 days after treating B16-F10 tumor bearing mice with one of the following: 1) no treatment controls (NT), 2) T cell only depots (TcellOnly_Depot), 3) SIVET with antigen-free FLT3L vaccine (SIVET_ FLT3L), 4) SIVET with antigen-free GMCSF vaccine (SIVET_ GMCSF), 5) antigen-free GMCSF vaccine-only depot (Vax_GMCSF) and 6) antigen-free FLT3L vaccine-only depot (Vax_ FLT3L) (Fig 3a). FMS-like tyrosine kinase 3 ligand (FLT3L)^26^ and Granulocyte-macrophage colony-stimulating factor (GMCSF)^16^ were used to recruit antigen presenting cells (APCs), while the TLR9 agonist, class B CpG, was included in all the SIVET and vaccine-only conditions to activate recruited APCs^18^. The vaccines were formulated without antigen, as dying cells from the tumors could serve as antigen source for the peritumorally injected depots. Functional analyses revealed the presence of tumor reactive host T cells in lymph nodes and spleens after antigen re-stimulation in vitro, with SIVET and vaccine-only conditions showing higher percentages of IFNG expressing host CD4+ and CD8+ T cells (Fig 3b-e). The proportions of tumor reactive T cells present in the lymph nodes and spleens were higher in the GMCSF than in FLT3L conditions.

**Fig. 3.**
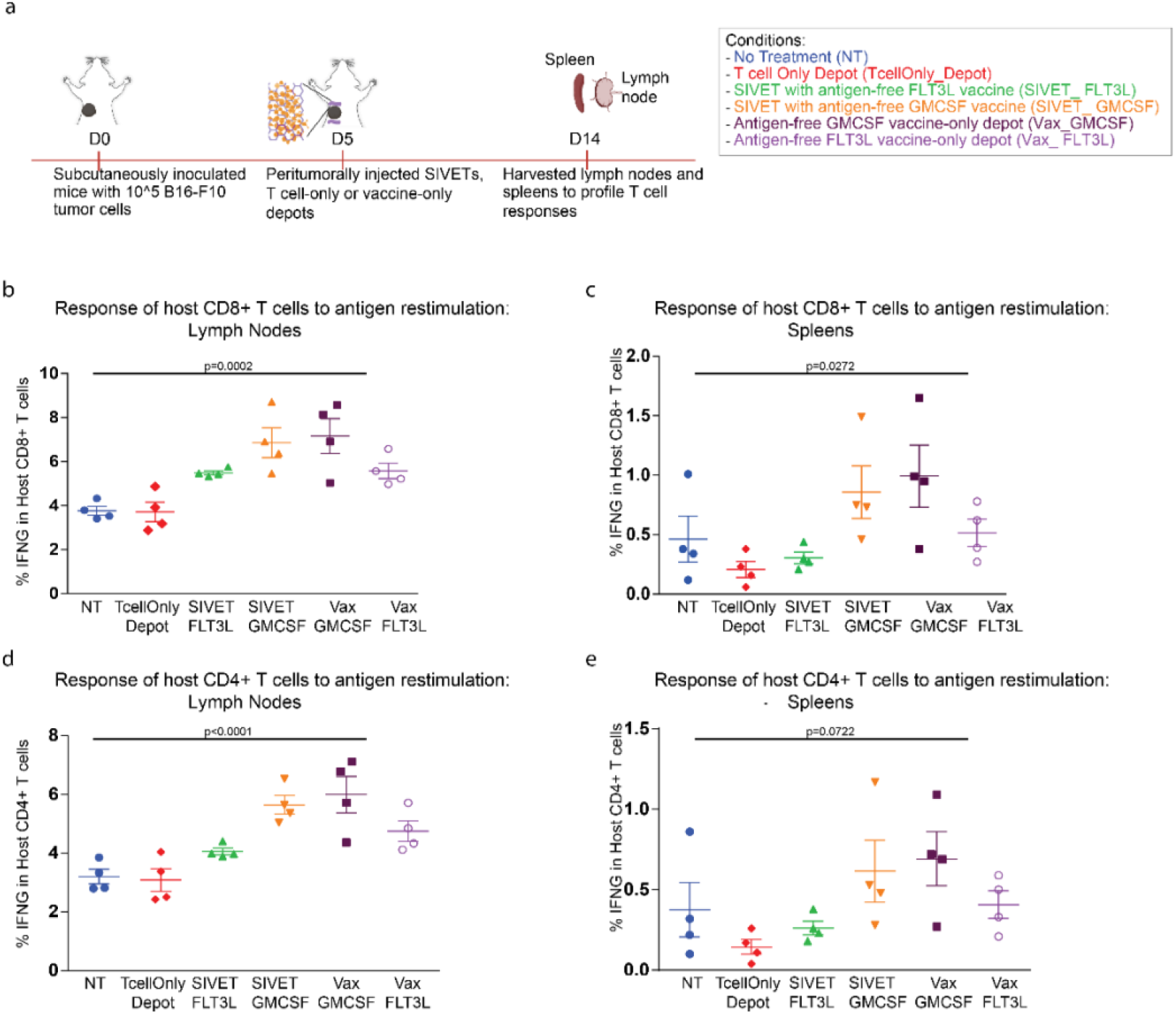
SIVETs elicit host T cell responses. a. Schematic of experiment. The following conditions were investigated: 1) no treatment controls (NT), 2) T cell only depots (TcellOnly_Depot), 3) SIVET with antigen-free FLT3L vaccine (SIVET_ FLT3L), 4) SIVET with antigen-free GMCSF vaccine (SIVET_ GMCSF), 5) antigen-free GMCSF vaccine-only depot (Vax_GMCSF) and 6) antigen-free FLT3L vaccine-only depot (Vax_ FLT3L). Proportions of IFNG expressing host CD8+ T cells isolated from lymph nodes (b) and spleens (c) after in vitro antigen re-stimulation. Proportions of IFNG expressing host CD4+ isolated from lymph nodes (d) and spleens (e) after in vitro antigen re-stimulation. p-values were determined by two-tailed one-way ANOVA with Geisser-Greenhouse correction. Data are mean ± s.e.m from n=4 mice per condition.

### H&E staining of both depots and tumor sections reveals treatment dependent cellular profiles

Hematoxylin and eosin (H&E) staining of both depots and tumor sections was next performed. Significantly higher cellularity was seen at the site of the SIVET_GMCSF depots relative to the other conditions, with many cells located at the interface between the depot and the native tissue, suggesting that the SIVETs do not trap the recruited cell populations. Importantly, there was markedly higher infiltration of immune cells at the tumor site for the SIVET treated conditions compared to the NT control, signaling a more pronounced immune activity in the former tumors (Supplementary Fig 3).

### SIVETs enhance the relative levels of activated antigen-presenting cells in depots and tumors

Broad phenotypic profiling of immune cells in both the tumors and depots was next performed to further understand the crosstalk between the adoptively transferred T cells and host immune cells (Fig 4a, Supplementary Fig 4a). The total numbers of immune cells infiltrating the tumors were influenced by their respective treatment condition, with SIVET_GMCSF having the highest levels of tumor infiltrating immune cells (Fig 4b). Accordingly, immunofluorescence imaging of myeloid cells in the tumors revealed a markedly high infiltration of myeloid cells in the SIVET_GMCSF condition compared with the TcellOnly_Depot and NT conditions (Fig 4c). Umap and Kmeans analyses also showed distinct immune cell populations within the tumor (Fig 4d-e, Fig 4f-h). Cluster 2, which comprises myeloid cells expressing canonical markers of activation and antigen presentation was enriched in the SIVET and vaccine-only conditions (Fig 4f-g). A CD11c-mid, CD11b-low, MHCII-low, CD80-low and CD86-low population (cluster 6), likely a non-activated myeloid population with poor antigen presentation, was most present in the NT control, with notable levels in the TcellOnly_Depot condition. Importantly, the SIVET conditions had lower proportions of cells in this non-activated myeloid population than the TcellOnly_Depot condition and their respective vaccine-only conditions (Fig 4g-h). Thus, SIVETs resulted in an enhancement of myeloid cells expressing markers of activation and antigen presentation and a concomitant reduction in non-activated myeloid populations with poor antigen presentation ability (Fig 4i).

**Fig. 4.**
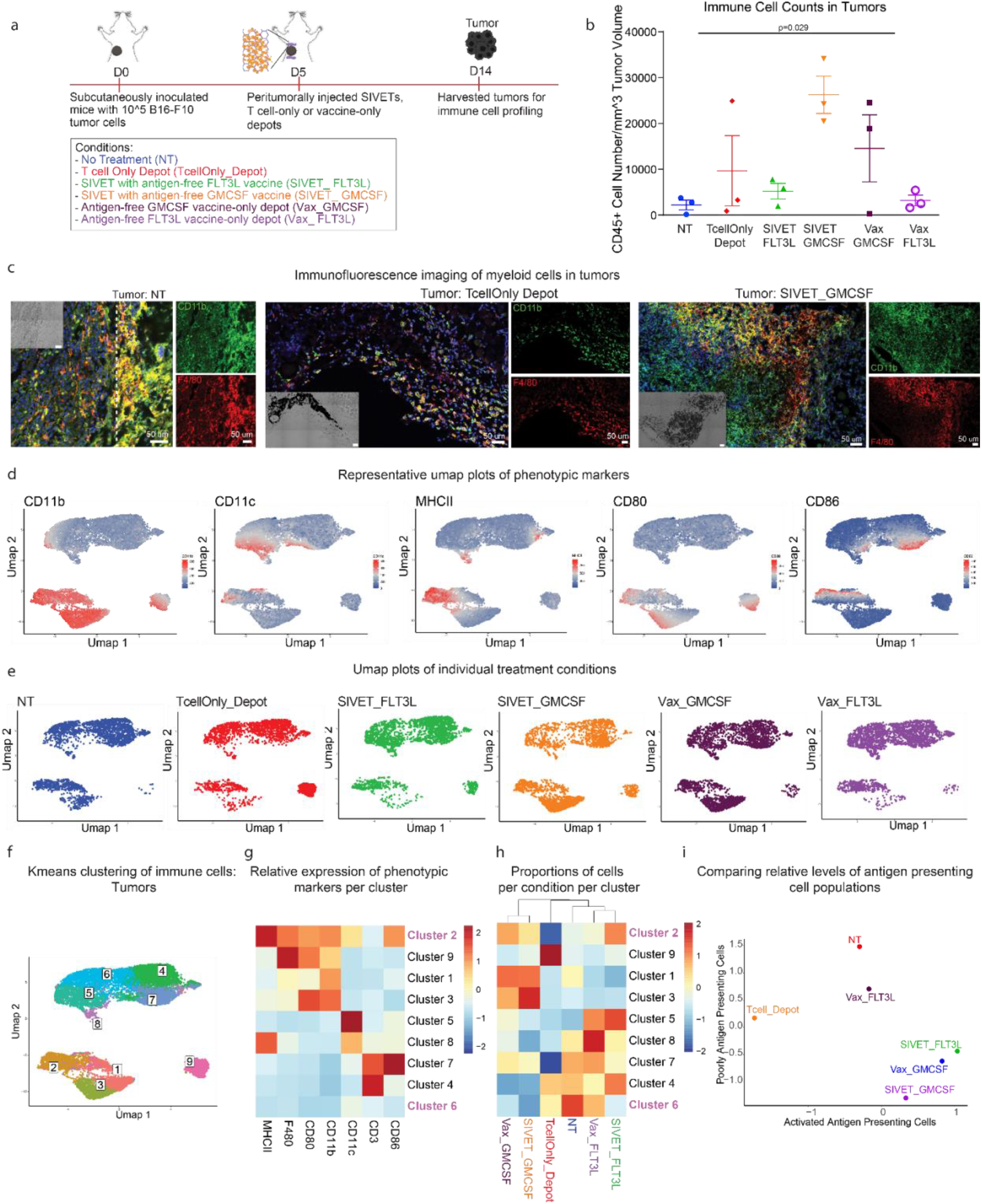
SIVETs enhance the relative levels of activated antigen-presenting cells in tumors. a. Schematic of experiment. b. Total numbers of tumor infiltrating CD45+ cells per mm^3 of tumor volume. p-value was determined by two-tailed one-way ANOVA with Geisser-Greenhouse correction. Data are mean ± s.e.m from n=3 mice per condition. c. Immunofluorescence imaging of myeloid cells expressing the indicated markers in NT, TcellOnly_Depot and SIVET_GMCSF tumors. d. Umap plots showing expression of the indicated markers. e. Umap plots of individual treatment conditions showing distinct localization of cells based on treatment group. f. Umap plot overlaid with Kmeans clusters of immune cells in tumors. g. Heatmap plot showing the average expression of the indicated immune cell markers in each cluster after K-means analysis. h. Heatmap plot showing the proportion of cells in each condition represented in each cluster. Some clusters of interest are highlighted. i. Scatterplot comparing the relative enrichment levels of the activated antigen presenting cell population (cluster 2), and the poorly antigen presenting cell population (cluster 6) as a function of treatment group. Data are pooled cells from n=3 mice per condition

The phenotypic profiles of immune cells in the depots mirrored those in the tumors. The total numbers of immune cells infiltrating the depots were dependent on treatment condition, with SIVET_GMCSF and Vax_GMCSF recruiting significantly more cells than the other conditions (Supplementary Fig 4b). Umap analyses of immune cells in the depots showed the presence of both T cell and myeloid cell populations (Supplementary Fig 4c). Immune cells in the various conditions were phenotypically distinct and localized to different regions on the umap plot (Supplementary Fig 4d). Cluster 5, which comprises a myeloid cell population expressing canonical markers of activation and antigen presentation, was most enriched in SIVET_ FLT3L and Vax_ FLT3L and least enriched in the TcellOnly_Depot condition (Supplementary Fig 4e-g). Importantly, the proportions of cells present in cluster 5 were higher in the SIVET conditions than their corresponding vaccine-only conditions, irrespective of the APC recruitment factor used (Supplementary Fig 4e-g). Additionally, when normalized for immune cell number, the proportions of the activated antigen presenting cells in cluster 5 was highest in the SIVET_GMCSF condition and remained higher in the SIVETs than their corresponding vaccine-only conditions (Supplementary Fig 4h).

### SIVETs minimize host T cell exhaustion in tumors and prolong activation of adoptively transferred T cells in depots

T cell specific profiles in tumors and depots were subsequently assessed (Fig 5a, Supplementary Fig 5a). T cell numbers infiltrating the tumors varied by treatment condition, with SIVET_GMCSF having the highest levels (Fig 5b). SIVET conditions had higher numbers of tumor infiltrating T cells than their respective vaccine only conditions (Fig 5b). Consequently, immunofluorescence imaging of T cells in tumors showed a noticeably higher T cell presence in the SIVET_GMCSF tumors relative to the NT control (Fig 5c). Both adoptively transferred, and host T cells were present in tumors of TcellOnly_Depot and SIVET_GMCSF treated groups (Fig 5d). Umap and Kmeans analyses of T cells in tumors also revealed distinct localization of T cell populations (Fig 5e-f). The adoptively transferred T cell population (cluster 1) was enriched in the TcellOnly_Depot condition tumors, while the SIVET conditions had higher proportions of host T cells (Fig 5e-i). Phenotypic analyses of host T cells in the tumors revealed a presumably exhausted CD8+ population expressing high levels of LAG3 and PD1 (cluster 3) (Fig 5e-i). Both umap and kmeans analyses showed that while this presumably exhausted T cell cluster (cluster 3) was mostly enriched in the NT control, the vaccine-only conditions had notable proportions of these T cells, which were dramatically diminished in their corresponding SIVET conditions (Fig 5e-i). A second PD1 expressing, LAG3-low host T cell cluster (cluster 5), which likely comprises non-exhausted T cells that are actively engaging tumor cells, was notably present in the SIVET conditions with a higher enrichment relative to the TcellOnly_Depot (Fig 5e-i). SIVETs therefore resulted in T cell populations that actively engaged tumor cells while minimizing their exhaustion (Fig 5j).

**Fig. 5.**
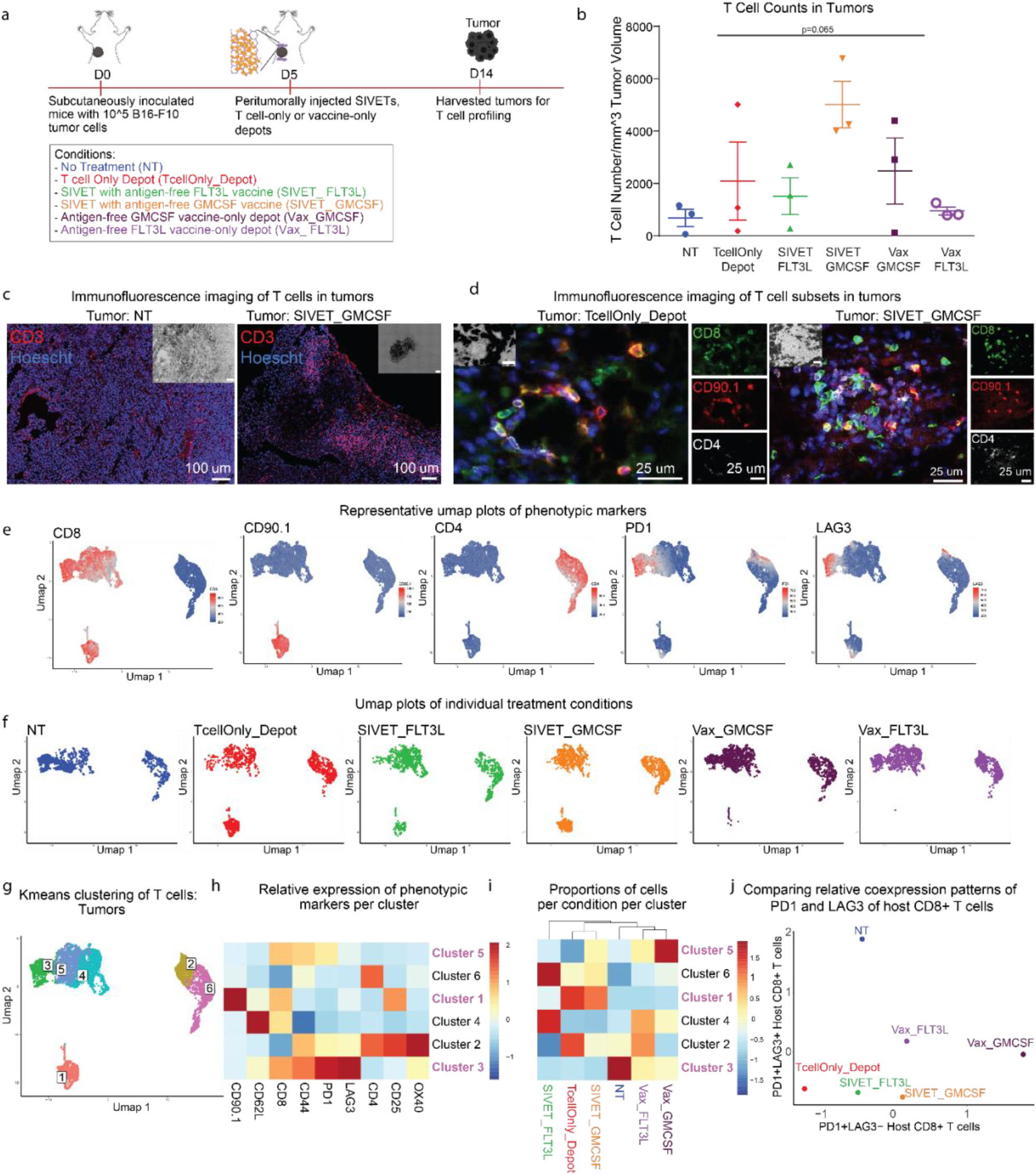
SIVETs minimize host T cell exhaustion in tumors. a. Schematic of experiment. b. Numbers of tumor infiltrating T cells per mm^3 of tumor volume. p-value was determined by two-tailed one-way ANOVA with Geisser-Greenhouse correction. Data are mean ± s.e.m from n=3 mice per condition. c-d. Immunofluorescence imaging of T cells present in NT and SIVET_GMCSF (c), as well as specific T cell subtypes in TcellOnly_Depot and SIVET_GMCSF tumors (d). e. Umap plots showing expression of the indicated markers. f. Umap plots of individual treatment conditions showing distinct localization of cells based on treatment group. g. Umap plot overlaid with Kmeans clusters of T cells in tumors. h. Heatmap plot showing the average expression of the indicated T cell markers in each cluster after K-means analysis. i. Heatmap plot showing the proportion of cells in each condition represented in each cluster. Some clusters of interest are highlighted. j. Scatterplot comparing the relative enrichment levels of PD1+LAG3-host CD8+ T cells (cluster 5), and PD1+LAG3+ host CD8+ T cells (cluster 3) as a function of treatment group. Data are pooled cells from n=3 mice per condition

Umap analyses of depots also showed treatment dependent effects on T cell profiles. Depots had both CD4+ and CD8+ T cells, with a significant CD90.1+ adoptively transferred T cell population present (Supplementary Fig 5b-c). The adoptively transferred T cells dominated the TcellOnly_Depot condition while the SIVET conditions had considerable infiltration of host T cells (Supplementary Fig 5b-c).

Kmeans clustering of the adoptively transferred T cells revealed two populations which had differential expression of CD25 and PD1. The SIVET conditions were enriched in the CD25-hi, PD1-mid population (cluster 5), while the TcellOnly_Depot condition was enriched in the CD25-low, PD1-low cluster (cluster 6) (Supplementary Fig 5d-f).

### SIVETs result in long-term tumor control

Using the B16-F10 tumor model, we next sought to ascertain whether the observed synergistic effects of SIVETs on both adoptively transferred T cell profiles and host immune cells translated into therapeutic benefits. Tumor growth analysis showed that while mice that received either therapeutic TcellOnly_Depot or vaccine-only treatments delayed tumor growth, they eventually succumbed to their tumors. However, mice treated with SIVETs significantly controlled tumor growth, with a majority of these mice completely rejecting their primary tumors. Combined survival analyses from 2 independent studies showed that 10/16 mice from SIVET_FLT3L and 13/16 mice from SIVET_GMCSF completely rejected their primary tumors long-term (Fig 6a-c).

**Fig. 6.**
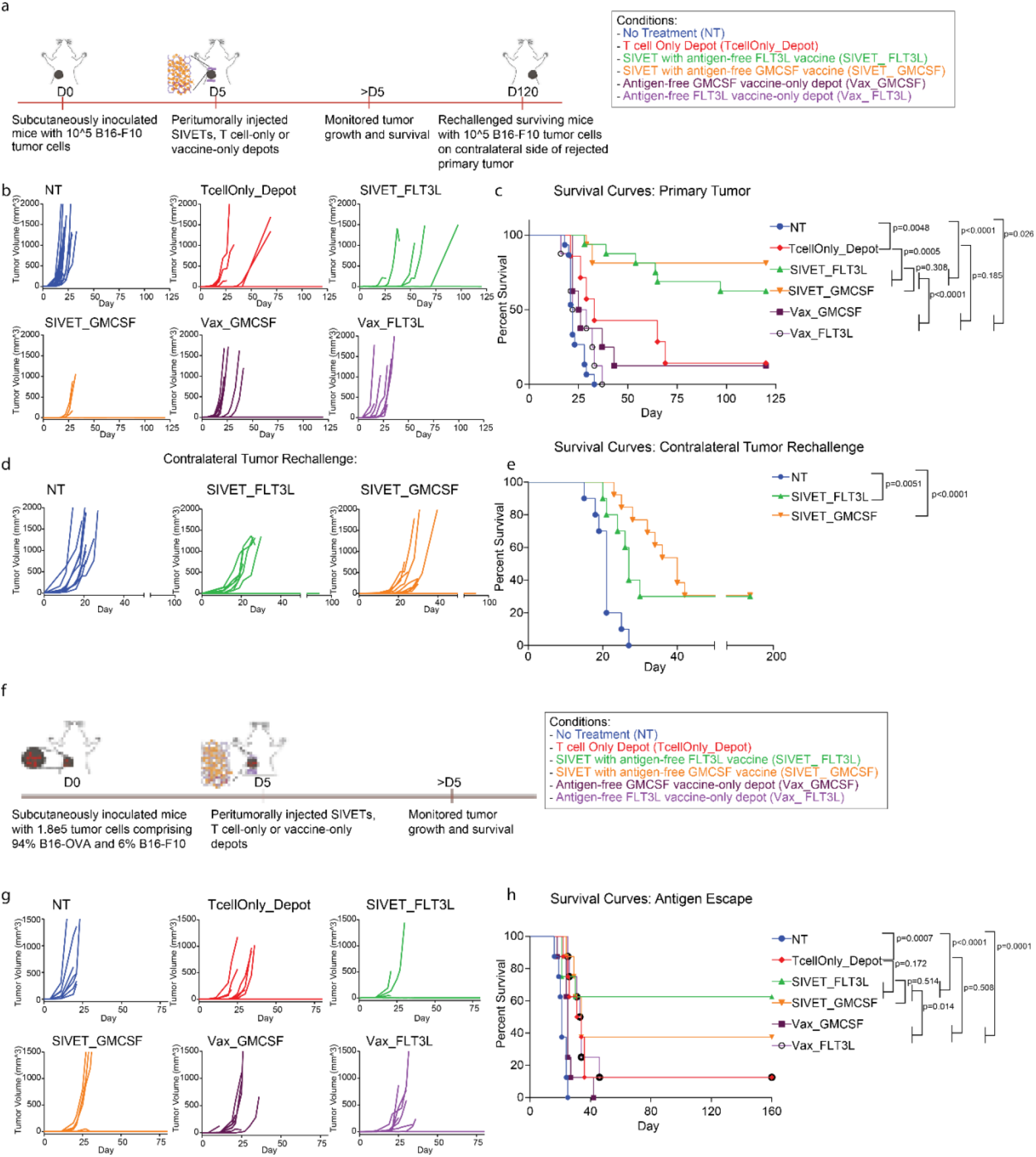
SIVETs enhance long-term tumor control. a. Schematic of experiment. b-c. Primary B16-F10 tumor studies. Tumor volumes (b) and Kaplan-Meier survival curves (c) comparing tumor growth and survival of mice treated with the indicated conditions. p-values for c were determined by Log-rank (Mantel-Cox) test. Data represent n=7-8 mice per condition for TcellOnly_Depot, Vax_FLT3L and Vax_GMCSF conditions and n=15-16 mice per condition, 2 independent studies (7-8 mice per study) for NT, SIVET_FLT3L and SIVET_GMCSF conditions. d-e. Contralateral tumor re-challenge studies for long-term surviving mice. d. Tumor growth (d) and Kaplan-Meier survival curves (e) of mice contralaterally re-challenged with 1e5 B16-F10 tumor cells after 120 days of primary tumor challenge. p-values for e were determined by Log-rank (Mantel-Cox) test. Data are n=10-13 mice per condition (2 independent studies). f-h. Antigen escape study. f. Schematic of therapeutic study for g-h. Tumor volumes (g) and Kaplan-Meier survival curves (h) comparing tumor growth and survival of mice treated with the indicated conditions. p-values for h were determined by Log-rank (Mantel-Cox) test. Data represent n=8 mice per condition.

Next, long-term surviving mice were re-challenged with the same dose of B16-F10 tumor cells contralateral to the location of the primary tumor after 120 days to assess the ability of memory T cells to mount systemic immune response against the secondary tumor. Tumor growth and survival analyses showed that long-term surviving mice from the SIVET_FLT3L and SIVET_GMCSF conditions significantly controlled their secondary contralateral tumors, with about a third of the mice in each condition completely rejecting their secondary tumors (Fig 6d-e). Interestingly, tumors grew slower in the SIVET_GMCSF condition compared with the SIVET_FLT3L condition for the mice that did succumb (Fig 6d-e).

### SIVET modulates long-term T cell phenotype

The impact of SIVET on long-term T cell phenotype was next investigated by profiling T cells at 82 days post tumor inoculation from the lymph nodes and spleens of mice that rejected their primary tumors. Only mice treated with SIVETs were profiled, in addition to naïve mouse controls, as the mice from the other conditions had succumbed to their tumors. Umap and Kmeans analyses of T cells in both lymph nodes and spleens showed that T cell profiles in the SIVET-treated mice were distinct from those of naïve mice (Supplementary Fig 6-7). In the lymph nodes, SIVET treatment resulted in an enrichment of more differentiated effector memory populations (CD62L-low, CD44-hi) for both CD8+ and CD4+ compartments (cluster 4 and cluster 8), compared with T cells from the naïve mice, which were mostly enriched in central memory (CD62L-hi, CD44-hi) and naïve (CD62L-hi, CD44-low) T cell clusters (clusters 5 and 6 respectively) (Supplementary Fig 6b-f). Importantly, umap, Kmeans and principal component analyses (PCA) revealed that SIVET_GMCSF was enriched in the more differentiated T cell clusters, while SIVET_FLT3L was intermediate (Supplementary Fig 6g). This trend was maintained in T cells harvested from spleens, where the SIVET_GMCSF condition was again most enriched in the more differentiated T cell clusters (Supplementary Fig 7).

Next, the long-term phenotype of the adoptively transferred CD90.1+ T cells was assessed. Adoptively transferred T cells were detectable in the lymph nodes and spleens of SIVET treated mice 82 days after tumor inoculation (Supplementary Fig 8b-c). Umap and Kmeans analyses of the adoptively transferred T cells from the lymph nodes revealed that the majority of the T cells were central memory (CD62L-hi, CD44-hi) T cells (clusters 1 and 5) (Supplementary Fig 8d-f). Importantly, there was a notable enrichment of SIVET_GMCSF in cluster 3, which is a more differentiated effector memory (CD62L-low, CD44-hi) T cell population (Supplementary Fig 8f-h), mirroring the observed phenotypic trends from the host T cell populations.

### SIVET provides enhanced protection against tumor antigen escape

Finally, we tested the ability of the SIVET-induced host T cell response to provide protection against tumor antigen escape by inoculating mice with a mixture of 94% B16-OVA tumor cells and 6% wild-type B16-F10 cells. B16-OVA and B16-F10 share common tumor antigens, but do not share the ovalbumin antigen. Mice were subsequently treated with OT1 CD8+ T cells, which recognize the ovalbumin peptide residues 257-264, and monitored for their ability to control both the B16-OVA and B16-F10 tumor cells. Tumor growth and mouse survival analyses showed that while the TcellOnly_Depot and the vaccine-only conditions delayed tumor growth and mouse survival, most of the mice eventually succumbed to the tumors. In contrast, about 60% and 40% of mice treated with either SIVET_FLT3L or SIVET_GMCSF, respectively, completely rejected their tumors long-term (Fig 6f-h).

## Discussion

Here we addressed the hypothesis that a biomaterial system which simultaneously enhanced adoptive T cell therapy and provided host immune engagement could provide synergistic immune protection, as evidenced by both immediate tumor debulking and long-term protection against solid tumors. While T cell depots by themselves showed superior therapeutic benefits relative to direct peritumoral T cell injection and IV T cell injection, SIVET resulted in synergistic effects between the adoptively transferred T cells and the host immune cells and led to long-term tumor control in most of the mice. Importantly, SIVET also provided enhanced protection against tumor antigen escape.

When used for localized T cell-only delivery, injectable T cell depots resulted in enhanced therapeutic benefits in solid tumors relative to IV and direct peritumoral T cell injection. These observations are consistent with previous studies that have reported greater therapeutic effects of biomaterial-based adoptive T cell delivery relative to traditional modes of T cell delivery^6–10^. While intravenous delivery could be optimal for hematological cancers, T cells delivered this way must traffic to solid tumors, and only a fraction of the adoptively transferred T cells find the tumor at a given time. T cells delivered peritumorally or intratumorally without scaffolds overcome the initial hurdle of trafficking, but generally do not persist at the tumor site, likely due to harsh conditions of the tumor microenvironment^27^.

Biomaterial-based delivery systems serve as T cell reservoirs that help to overcome the hurdle of trafficking while providing the requisite environment for T cell persistence. Previous biomaterial-based platforms used for adoptive T cell delivery have mostly been in the form of spherical or sheet-like implantable scaffolds^6–8^, or injectable in-situ forming gels^9,10^. Implantable scaffolds require invasive surgeries for administration, and the spherical or sheet-like shapes could pose difficulties in terms of scaling for larger mammals. While injectable, in-situ forming gels may be effective in anatomical locations where an injection pocket could be formed (eg. subcutaneous space), they likely would have limited applicability in other locations due to dispersion from the injection site before gelation is complete. The injectable, rod-shaped T cell depot described in this study overcomes these challenges, as it obviates the need for invasive surgery and allows for the use of syringes and catheters with minimal shear. The length of the depots can be easily tuned for larger mammals. The pre-formed depots could also be injected at multiple anatomical locations while allowing the depots to conform to the area of injection, because of their flexibility.

SIVET resulted in synergistic phenotypic effects between the adoptively transferred T cells and host immune cells. The recruitment and activation of host antigen presenting cells correlated with greater activation of the adoptively transferred T cells, relative to the T cell only depot condition. This is consistent with reports that continual help from antigen presenting cells is important for anti-tumor efficacy of transferred T cells^28^. Further, it is likely that the reduction in the proportions of non-activated, poorly antigen presenting myeloid cells in the SIVETs compared with vaccine-only conditions is due to paracrine signaling from the adoptively transferred T cells to the recruited myeloid cells. The dramatic decrease in host CD8+ T cell exhaustion in SIVET treated tumors relative to the vaccine-only conditions could be due to tumor debulking by the adoptively transferred T cells that reduces antigen density, limiting over-stimulation of intra-tumoral host T cells. The specific depot agent (GM-CSF vs FLT3L) utilized to concentrate host myeloid cells informed the composition of myeloid cells at the depot and tumor sites, and influenced the long-term phenotypic profiles of both the adoptively transferred and host T cells. FLT3L particularly recruited classical antigen presenting myeloid cell populations, while GMCSF recruited a broader array of myeloid populations and in higher numbers, consistent with the expected behavior of the two cytokines^29,30^. Additionally, the availability of T cell populations with faster effector response, as seen in the higher proportions of long-term effector memory adoptively transferred and host T cell populations for the SIVET_GMCSF condition, could explain why tumors grew slower in SIVET_GMCSF than SIVET_FLT3L upon secondary challenge for mice that succumbed.

SIVET provided enhanced long-term tumor control and protection against tumor antigen escape, confirming that the synergistic phenotypic effects of SIVET on both adoptively transferred T cell profiles and host immune cells translated into long-term therapeutic benefits. Previous studies have shown that approaches that promote resistance of CAR T cell to exhaustion^31^ or directly boost their expansion in vivo^32^ can lead to better therapeutic responses to solid tumors. Enhanced efficacy of adoptive T cell therapy can also be promoted by engaging host immunity with cGAMP^7^. Our study demonstrates phenotypic and therapeutic synergy between adoptive T cell therapy and the active recruitment of antigen presenting cells for in-situ vaccination, utilizing principles from previous in-situ vaccination work^20,33–37^. The enhanced activation of the adoptively transferred T cells in the SIVET conditions as well as the dramatic decrease in host T cell exhaustion likely allow both adoptively transferred and host T cells to provide enhanced anti-tumor protection individually, and to synergize to provide long-term therapeutic benefits against aggressive tumors.

In sum, we have demonstrated a strategy to enhance adoptive T cell therapy with therapeutic cancer vaccination to achieve long-term anti-tumor efficacy in murine solid tumors, providing a potential path to overcoming some of the clinical limitations of adoptive T cell therapy in this setting. These findings further motivate the development of additional strategies that can simultaneously provide local and enhanced adoptive T cell therapy, while eliciting host T cell responses.

## Methods

### Material synthesis

#### Collagen modification

Rat Tail Collagen Type I (Corning #354236) was modified with 5-Norbornene-2-acetic acid succinimidyl ester (Nb-NHS) (Sigma Aldrich #776173) using a ratio of 1g Nb-NHS: 10g collagen. First, rat tail collagen was neutralized with NaOH to pH 7.2-7.5, buffered with 10x DPBS and diluted to an initial concentration of 2mg/ml. Next, Nb-NHS was dissolved to 2mg/ml in DMSO and diluted 10x in 1x PBS. An equal volume of Nb-NHS solution was then added to the neutralized collagen under continual stirring, resulting in 1mg/ml final collagen concentration. The reaction proceeded for 5 hours at 4°C to delay collagen gelation and quenched with 0.1N acetic acid to re-acidify the collagen solution. Norbornene-modified collagen (Col-Nb) was then dialyzed against 0.025 N acetic acid for 4 days, filtered through a 0.45um filter and lyophilized.

#### Alginate modification

Alginate tetrazine was synthesized as previously described. Briefly, Pronova ultrapure MVG sodium alginate (Novamatrix) was modified with (4-(1,2,4,5-tetrazin-3-yl)phenyl methanamine hydrochloride (Karebay Biochem) using carbodiimide chemistry. Alginate was first dissolved in 0.1M 2-(N-morpholino)ethanesulfonic acid (MES), 0.3 M NaCl, pH 6.5 at 5 mg/ml. Next, 1.9g of ethyl-3-(3-dimethylaminopropyl)-carbodiimide hydrochloride (EDC) (ThermoFisher #22980) and 1.2g of N-hydroxysuccinimide (NHS) (ThermoFisher #24500) were added per gram of alginate. 0.1g of tetrazine was then added under constant stirring and allowed to react overnight at room temperature. The tetrazine modified alginate was subsequently purified first by tangential flow filtration (KrosFlow KR2i; Spectrum Labs) against a 150mM to 0mM decreasing NaCl gradient with a 1kDa MWCO membrane, and then by treatment with 1g activated charcoal. This was followed by filtration through a 0.22um filter and subsequent lyophilization.

### Fabrication of depots

To fabricate 0.75% wt/vol hybrid alginate-collagen cryogel depots comprising 60% alginate and 40% collagen, lyophilized Col-Nb was first dissolved to 9 mg/ml in 0.025 N acetic acid at 4°C for at least 48 hours. Alginate tetrazine was freshly dissolved on the day of cryogel fabrication to 2% wt/vol and cooled at 4°C. Dissolved Col-Nb was neutralized with cold 1N NaOH, buffered with cold 10x DPBS and balanced with cold milliQ water, after which the alginate tetrazine was added, yielding 3mg/ml and 4.5mg/ml final concentrations of collagen and alginate respectively. The cryogel mixture was immediately pipetted into a 2mm diameter tygon tubing (VWR #89404-318) at 50ul cryogel mixture per 1cm tubing, and placed in a −15°C freezer overnight for cryo-polymerization. After cryogelation, cryogels were thawed at room temperature and ejected by gently flushing the tygon tubing with 400ul DPBS. Depots with different ratios of alginate to collagen or total polymer content were fabricated by varying the relative final concentrations of the alginate or collagen (while keeping the total polymer amount constant), or by increasing the total amount of polymer respectively.

### Estimation of cryogel interconnected porosity and shape recovery

Cryogel interconnected porosity and shape recovery were estimated as previously described. Briefly, Cryogel interconnected porosity was investigated by first weighing intact cryogels *(M*_*hydrated*_), wicking cryogels with kimwipe for 15 seconds to remove excess buffer from cryogel pores and re-weighing the wicked cryogels *(M*_*wicked*_). Interconnected porosity was quantified as follows:

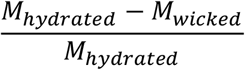

Cryogel shape recovery/memory was estimated by first weighing intact cryogels *(M*_*hydrated*_), wicking cryogels with kimwipe for 15 seconds to remove excess buffer, rehydrating the gels in DPBS and re-weighing the rehydrated cryogels *(M*_*rehydrated*_). Shape recovery was quantified as follows:

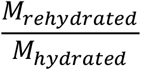

### SEM and SHG characterization of depots

#### SEM

Cryogels were fixed with 2.5% glutaraldehyde and 2% PFA in the presence of 0.115M sucrose and 100mM Hepes for 1 hour. Scaffolds were then washed in 100mM Hepes and 0.115M sucrose 3 times, each for 5 minutes, followed by 1% osmium staining for 1 hour and 1% Uranyl acetate staining overnight. After staining, samples were washed 3 times with milliQ water and serially dehydrated with 50%, 70%, 95% and 100% ethanol for 10 minutes each. Scaffolds were then freeze fractured in liquid nitrogen, dried using critical point drying, mounted on to stubs, coated with platinum/palladium for 10 seconds and imaged with the Ultra plus FESEM at 3keV with gun vacuum at 2.29E-10 mbar and system vacuum at 1.43E-06 mbar. Images were taken using the secondary electron (SE2) detector.

#### SHG

Pristine cryogels were imaged via second harmonic generation using a Leica SP5 X MP Inverted Confocal Microscope at 820nm wavelength at 10x magnification. Pore size distribution was determined using Imaris.

### Incorporation of bioactive factors into depots

Bioactive factors like IL2, GMCSF, FLT3L and CpG were incorporated into cryogels by adding them to the gel mixture before cryogelation. To achieve controlled release, recombinant mouse IL2 (Biolegend #575408) and recombinant murine GMCSF (Peprotech #315-03) or recombinant murine FLT3L (Peprotech #250-31L) were preloaded onto charged laponite XLG (BYK additives) as follows. First, laponite was dissolved at 30mg/ml in milliQ with extensive vortexing until solution became clear. For cryogels loaded with both IL2 and GMCSF/FLT3L, 5ug of IL2/50ul gel solution and 1ug of GMCSF or FLT3L/50ul gel solution were each separately adsorbed onto 25ug laponite for 1 hr at 4°C. For cryogels loaded with IL2 only, 5ug of IL2/50ul gel solution was adsorbed onto 50ug laponite for 1 hr at 4°C. This kept the total amount of laponite in the scaffold at 50ug. 50ug of CpG ODN 1826 (Invivogen #tlrl-1826-1) was adsorbed onto 3ug linear 25K PEI (Polysciences 23966) for 1 hr at 4°C. The preloaded bioactive factors were then added to the gel mix before gels underwent cryogelation as described above.

CpG release studies was performed using the Quant-iT OliGreen ssDNA Kit (Thermo #O11492), while cytokine release was quantified via ELISA.

### T cell isolation, activation, culturing

Splenic mouse CD8+ T cells were isolated with magnetic-bead-based CD8a+ T Cell Isolation Kit, mouse (Miltenyi #130-104-075) following the manufacturer’s protocol. T cells were activated with Dynabeads® Mouse T-Activator CD3/CD28 (ThermoFisher Scientific # 11452D) at 1:1 ratio in T cell media: RPMI 1640 (Lonza #BE12-702F), 10% heat-inactivated fetal bovine serum (Gibco #10-082-147), 1% pen/strep, 55 μM *β*-mercaptoethanol, 10 mM HEPES (Sigma Aldrich # H4034), 1% 100x non-essential amino acid (Lonza #13-144E), 100 mM sodium pyruvate (Lonza #13-115E), supplemented with recombinant mouse IL-2 (BioLegend #575404).

### T cell Migration Assay

To perform T cell migration, T cells were first stained with 0.5uM cell tracker deep red (ThermoFisher #C34565) and loaded into pre-formed cryogels by briefly wicking the scaffolds using kimwipe and rehydrating them in 25ul of 2e5 cells/ml concentrated T cell solution. Time-lapse imaging was taken at 20x magnification for 2 hrs at 60s intervals and analyzed using Imaris.

### Animal Studies

Animal studies were performed in accordance with the National Institutes of Health and the Harvard University Faculty of Arts and Sciences’ Institutional Animal Care and Use Committee (IACUC) guidelines.

#### In vivo T cell persistence study

To investigate local in vivo T cell persistence following administration of T cell loaded depots, CD8+ T cells were first isolated from the spleens of luciferase expressing female FVB-Tg(CAG-luc,-GFP)L2G85Chco/J mice (Jackson #008450) and activated for 4 days as described above, after which the dynabeads were removed. Depots loaded with 5ug recombinant murine IL2 adsorbed onto 50ug laponite were infused with 2e6 luciferase expressing CD8+ T cells. Two depots per mouse were subcutaneously injected into the left flank of female C57 albino mice (Jackson #000058). 4e6 luciferase expressing CD8+ T cells with matched IL2 were directly injected subcutaneously to serve as controls. Luciferase expression was tracked using bioluminescence imaging by subcutaneously administering 150mg/kg of D-luciferin (GoldBio #LUCK-100) and measuring luminescence via IVIS (Perkin Elmer)

#### B16-F10 therapeutic tumor studies

*Tumor inoculation:* 1e5 B16-F10 tumor cells at passage 7 (ATCC #CRL-6475) were subcutaneously injected into the left flank of 7-9 weeks old female C57BL/6J mice (Jackson #000664) on day 0 and treated on day 5, when the tumors had become palpable. For studies with sublethal irradiation preconditioning, mice were subjected to 5Gy gamma irradiation on day 4. *T cell isolation and loading into depots:* CD8+ T cells were isolated from the spleens of B6.Cg-Thy1a/Cy Tg(TcraTcrb)8Rest/J (pmel) mice (Jackson #005023), which recognize the gp100 antigen on B16-F10 tumor cells, and activated using dynabeads for 4 days. The activated T cells were then separated from the dynabeads and infused into depots at 2e6 T cells/cryogel. *Depot composition:* Depots were fabricated as detailed above. Depots designated for T cell delivery only were loaded with 5ug of recombinant murine IL2 adsorbed onto 50ug laponite. SIVETSs had 5ug recombinant murine IL2 and 1ug recombinant murine GMCSF or FLT3L individually adsorbed onto 25ug laponite, as well as 50ug CpG ODN 1826 adsorbed onto PEI. Vaccine only depots received the same formulation but without T cell infusion.

*Injection of depots:* Two depots were injected per mouse, peritumorally on either side of the tumor in 100ul DPBS using a 16-gauge needle. For IV T cell injection and direct peritumoral injection, 4e6 pmel T cells with matched IL2 were administered either by tail vein injection or injected subcutaneously next to the tumor in 100ul DPBS using a 25-gauge needle. *Tumor volume measurements and mice survival:* For tumor volume determination, calipers were used to measure the tumor length, width and height. Tumor volume was estimated as 0.5 ∗ *length* ∗ *width* ∗ *height*. Where the tumor was flat, a width of 0.5mm was assumed. Mice were euthanized when tumors grew to 20mm in any dimension, when tumors became necrotic or when excessive weight loss was observed in tumor bearing mice.

#### Antigen escape tumor model

1.8e5 tumor cells comprising 94% B16-cOVA (a kind gift from the Wucherpfennig lab) and 6% B16-F10, both at passage 7 were subcutaneously injected to the into left flank of 8 weeks old C57BL/6J mice. CD8+ T cells were isolated from the spleens of female C57BL/6-Tg(TcraTcrb)1100Mjb/J (OTI) mice (Jackson #003831), which recognize the ovalbumin peptide residues 257-264, and activated using dynabeads for 4 days. T cells were separated from the dynabeads and infused into the depots as described above. Depot composition, tumor volume measurements and criteria for mice survival are the same as previously described.

### Tissue processing

#### Tumors

Tumors were excised into gentleMacs C tubes (Miltenyi #130-093-237) containing 150U/ml collagenase type IV (Thermo #17104019) and 0.1ug/ul DNAse 1 (Sigma #11284932001) in digestion media: RPMI 1640+10% FBS. Tumors were mechanically dissociated using the gentleMacs tissue dissociator (Miltenyi #130-093-235) program m_spleen_03, and incubated for 25 minutes at 37°C. Tumors were then mechanically dissociated for the second time using the same program and incubated for 15 extra minutes. The enzymatic reaction was quenched using MACS buffer: DPBS with 0.5% BSA and 2mM EDTA, and filtered through a 30um strainer (Miltenyi #130-098-458)

#### Lymph nodes

Brachial, axillary and inguinal lymph nodes were harvested and pooled before digestion. The same digestion protocol highlighted above was used to digest the lymph nodes.

#### Depots

Depots were harvested into gentleMacs C tubes containing 150U/ml collagenase type IV, 0.1ug/ul DNAse 1 and 2U/ml alginate lyase (Sigma #A1603). The same digestion protocol described above was used to digest depots.

#### Spleens

Harvested spleens were mechanically dissociated by using 1ml syringe plungers to mash the spleens through 30um strainers. Red blood cell lysis (BioLegend #420302) was performed on single cell suspension for 1 min before further downstream processing.

### In vitro T cell restimulation

Lymph nodes and spleens were first digested as described above. To perform in vitro T cell stimulation, lymph node and spleen derived cells were incubated with a cocktail of 2ug/ml mgp100, M27 and M30 peptides in addition to 5e4 B16-F10 tumor cells. The broad approach to antigen stimulation was taken because the vaccine was antigen-free, and thus a broad repertoire of T cell clones was expected to be elicited by the vaccines. After 1.5hrs, 0.27ul of the GolgiStop protein transport inhibitor (BD #554724) was added to each well, after which the cells were incubated for 4 hrs. The cells were then processed for flow cytometry.

### Flow cytometery

#### Surface staining

Cells were kept at 4°C throughout immunostaining. First, cells were stained with LIVE/DEAD™ Fixable Blue Dead Cell Stain (ThermoFisher Scientific #L23105) at 1000x dilution for 30mins in PBS, after which staining was quenched with flow cytometry staining (FACs) buffer (Invitrogen #00-4222-26). Cells were blocked with TruStain FcX Fc receptor blocking solution (BioLegend #101319) for 5 min and stained with surface protein antibodies for 20 min, after which the cells were washed 3x in FACs buffer. Flow cytometry acquisition was performed on a BD Fortessa LSRII. Single color compensation beads (Thermo #01-2222-41), was used for multi-parameter flow cytometry compensation. Gating was done based on fluorescence-minus-one (FMO) controls.

#### Intracellular cytokine staining (ICS)

ICS was performed after live/dead and surface staining, using the Cyto-Fast™ Fix/Perm Buffer Set (BioLegend #426803) according to the manufacturer’s protocol.

Briefly, cells were fixed in the Cyto-Fast™ Fix/Perm Buffer for 20 minutes at room temperature, washed twice in 1X Cyto-Fast™ Perm Wash solution and stained in 1X Cyto-Fast™ Perm Wash solution for 20 minutes at room temperature. After staining, cells were washed 3x in FACs buffer before acquisition on the BD Fortessa LSRII.

### Flow cytometry analyses

#### Flowjo analyses

Fcs files exported from the BD Fortessa LSRII cytometer were imported into Flowjo, analyzed using the following hierarchy: SSC-A/FSC-A to gate for lymphocytes → FSC-H/FSC-A FSC-H/FSC-A to gate for single cells → CD3/Live_Dead to gate for live T cells (or CD45/Live_Dead for total immune cells) → CD4/CD8 to gate for CD4 or CD8 T cells. Further downstream gating is performed using FMO controls, or compensated single cell flow cytometry intensity values are exported as csv files for unsupervised analyses. Sample gating strategy is shown in Extended Fig 11

#### Unsupervised analyses

*Umap analyses:* Exported single cell flow cytometry intensities were imported into R version 4.0.5 and pooled together. Outlier intensities, defined as values outside 3 standard deviations from the mean (in both directions), were discarded. Cells were then down sampled to keep the same number of cells per condition, after which umap analyses were performed. Umap projections were plotted as 2D scatter plots using ggplot2 version 3.3.3^38^, and annotated with either the marker intensity or condition. Umaps of the individual conditions were plotted either as 2D scatter plots or as 2D scatter heatmaps (LSD R package 4.1.0^39^) by subsetting the umap projections of the condition of interest from the projection of the pooled dataset. *Kmeans:* Kmeans was performed by first discarding outlier intensities as described above. To estimate the appropriate number of clusters (K), and Elbow plot was generated by estimating the total within sum of squares iterated over K=1 to K=20. The optimum K was chosen as the Elbow point on the plot, where increasing K does not lead to substantial changes in the total within sum of squares. Using the optimum number of clusters, Kmeans clustering was performed on the cells, after which the cluster each cell belongs to was overlaid onto the independently generated umap plot to show concordance. To determine the phenotypic markers that characterize a specific cluster, the cluster centers, which represent the mean expression of each marker in each cluster were estimated. The proportions of cells per condition per cluster were also determined and were represented as heatmaps using the Pheatmap package version 1.0.12^40^. *Principal component analysis (PCA):* PCA was performed on the frequencies of each condition in each cluster to assess the similarity between the different conditions. PCA scores and loadings for the first two principal components were then plotted using the factoextra package version 1.0.7^41^ in R.

### Immunofluorescence imaging

Tumors and were excised and fixed for 1hr in 4% PFA at 4°C. Tissues were cryopreserved in 30% sucrose overnight and then frozen in OCT solution. Tissues were then cryo-sectioned into 20-30 µm slices and mounted on superfrost plus slides. For immunostaining, the OCT was removed by immersing the sections in 1xPBS, blocked with 5% normal rat serum, 5% normal mouse serum, and purified rat anti-mouse CD16/CD32 for 30 min. Finally, samples were stained using the manufacturer’s specifications.

Slides were washed 2x with 1xPBS, stained with Hoescht 33342, mounted in ProLong Gold Antifade Mountant (Thermo Fisher) and covered with a no. 1.5 coverslip (Electron microscopy services). Tissues were imaged via tiled acquisitions on an LSM 710 confocal microscope using a 32x water immersion objective. Images were stitched and processed in the Zen Black software.

### H&E

Tissue sections were manually stained by H&E. Briefly, sections were hydrated using decreasing concentrations of ethanol, and then stained by hematoxylin (Mayer’s hemalum solution, Sigma, USA). Differentiation and bluing agents (Differentiation solution and Scott’s Water, both from Sigma, USA) were then used to augment hematoxylin staining. Eosin stain (Eosin Y solution, Sigma USA) was then applied, followed by ethanol dehydration. Imaging was done on an Echo Revolve Microscope.

### Statistical Analyses

Unless otherwise specified, all statistical analyses were performed on Prism Graphpad software version 9.0.2. Statistical tests used Student’s t-test with Welch’s correction for comparison between two groups and two-tail one-way ANOVA with Geisser-Greenhouse correction for group comparisons. Two-way ANOVA with repeated measures were used for the release study datasets and Log-rank (Mantel-Cox) test for mouse survival studies. P-value less than 0.05 was considered as significant unless otherwise noted.

Error bars represent standard error of mean, unless otherwise noted.

## Supplementary Figures

**Supplementary Fig. 1.**
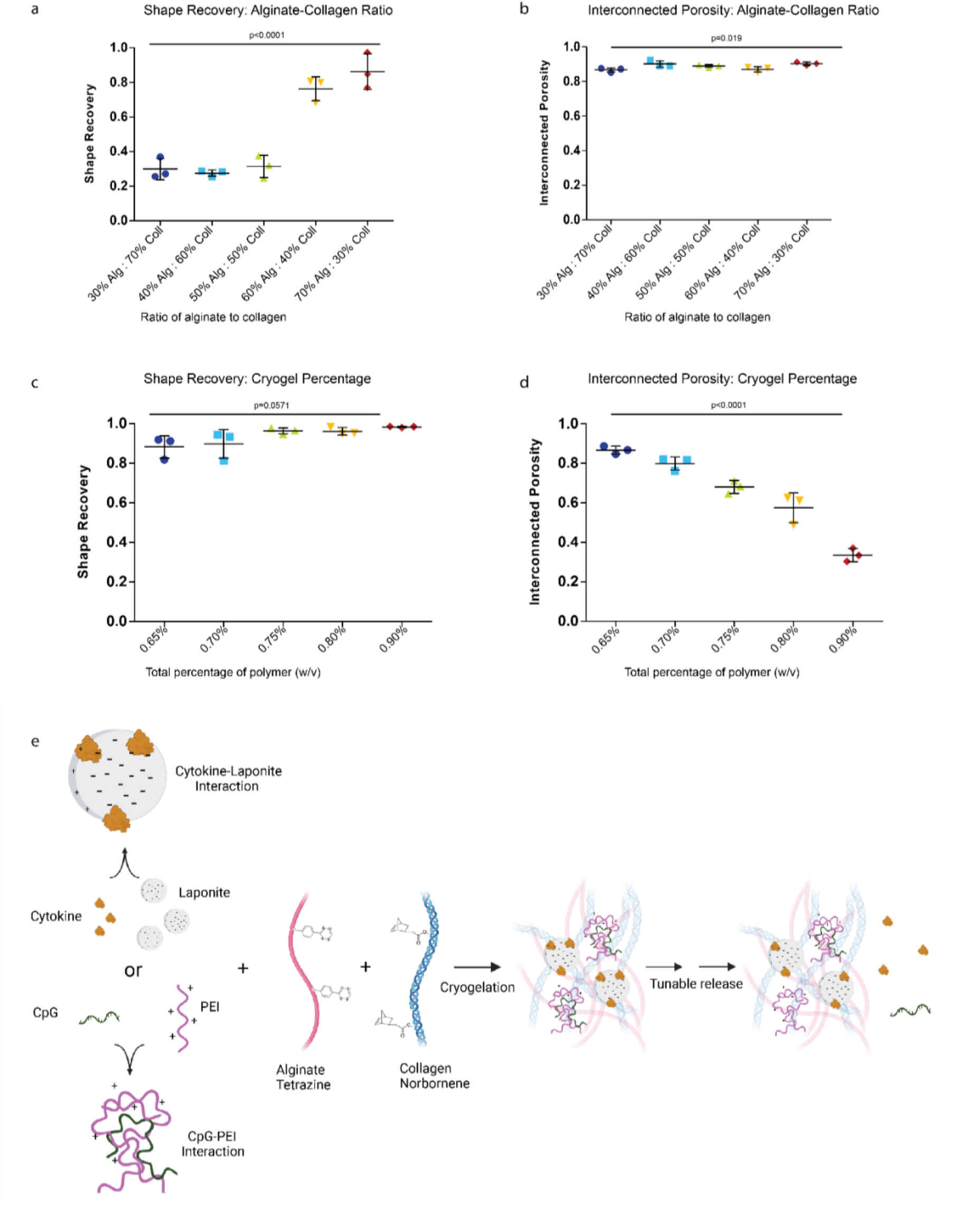
Further characterization of SIVETs. a-b. Cryogel shape recovery and interconnected porosity profiles as a function of changing the ratios of alginate to collagen. c-d. Same profiles as in a-b as a function of varying total percentage of polymer. P-values determined by two-tailed one-way ANOVA with Geisser-Greenhouse correction. Data represent mean ± s.e.m from n=3. c. Schematic of tunable release of bioactive factors from depots. To achieve tunable release of bioactive factors, cytokines (eg. IL2, GMCSF) and adjuvants (CpG) were either directly added to cryogel mixture before cryogelation, or pre-adsorbed onto laponite (for cytokines) or PEI (for CpG) before they were added to cryogel solution.

**Supplementary Fig. 2.**
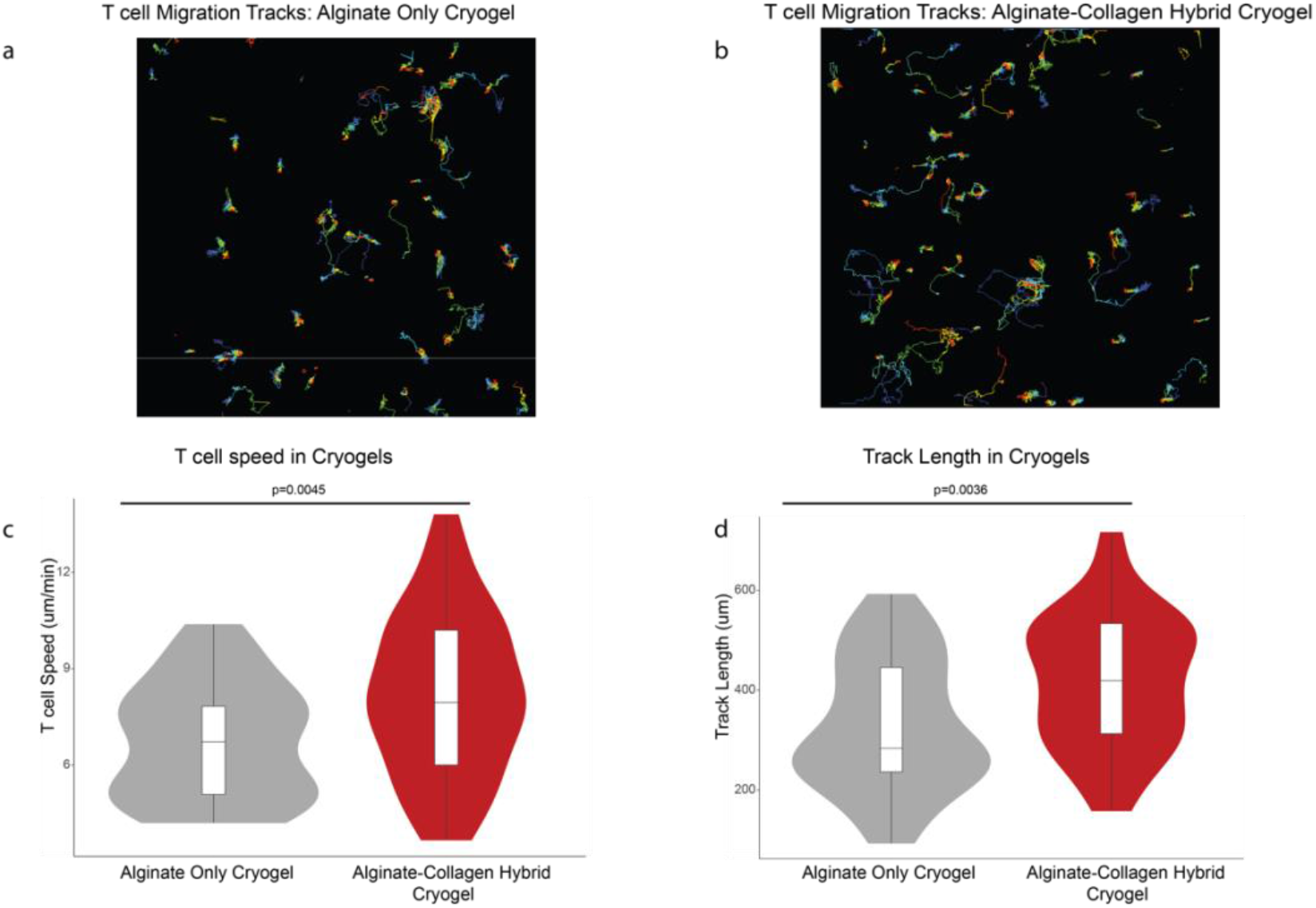
Collagen in depots enhances T cell migration in vitro. a-b. Migration tracks for T cells loaded in alginate-only cryogels (a) or alginate-collagen hybrid depots (b). c-d. Violin plots comparing T cells speeds (c) and track lengths (d) for T cells loaded in alginate and alginate-collagen hybrid depots. P-values determined by two-tailed unpaired t test with Welch’s correction.

**Supplementary Fig. 3.**
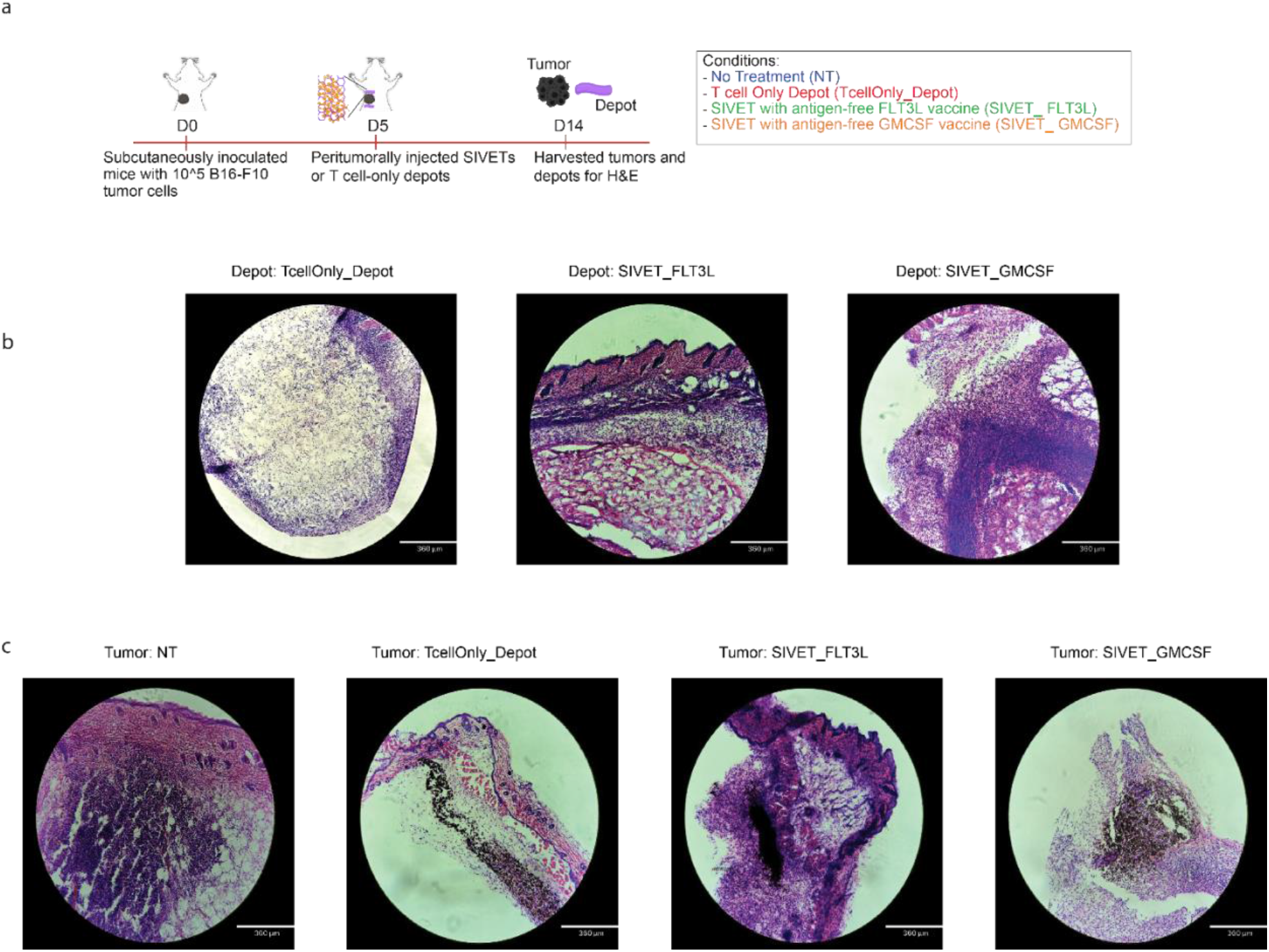
H&E images of depots and tumors after SIVET treatment. a. Schematic of experiment. Representative H&E images of depots (b) and tumors (c) for the indicated conditions.

**Supplementary Fig. 4.**
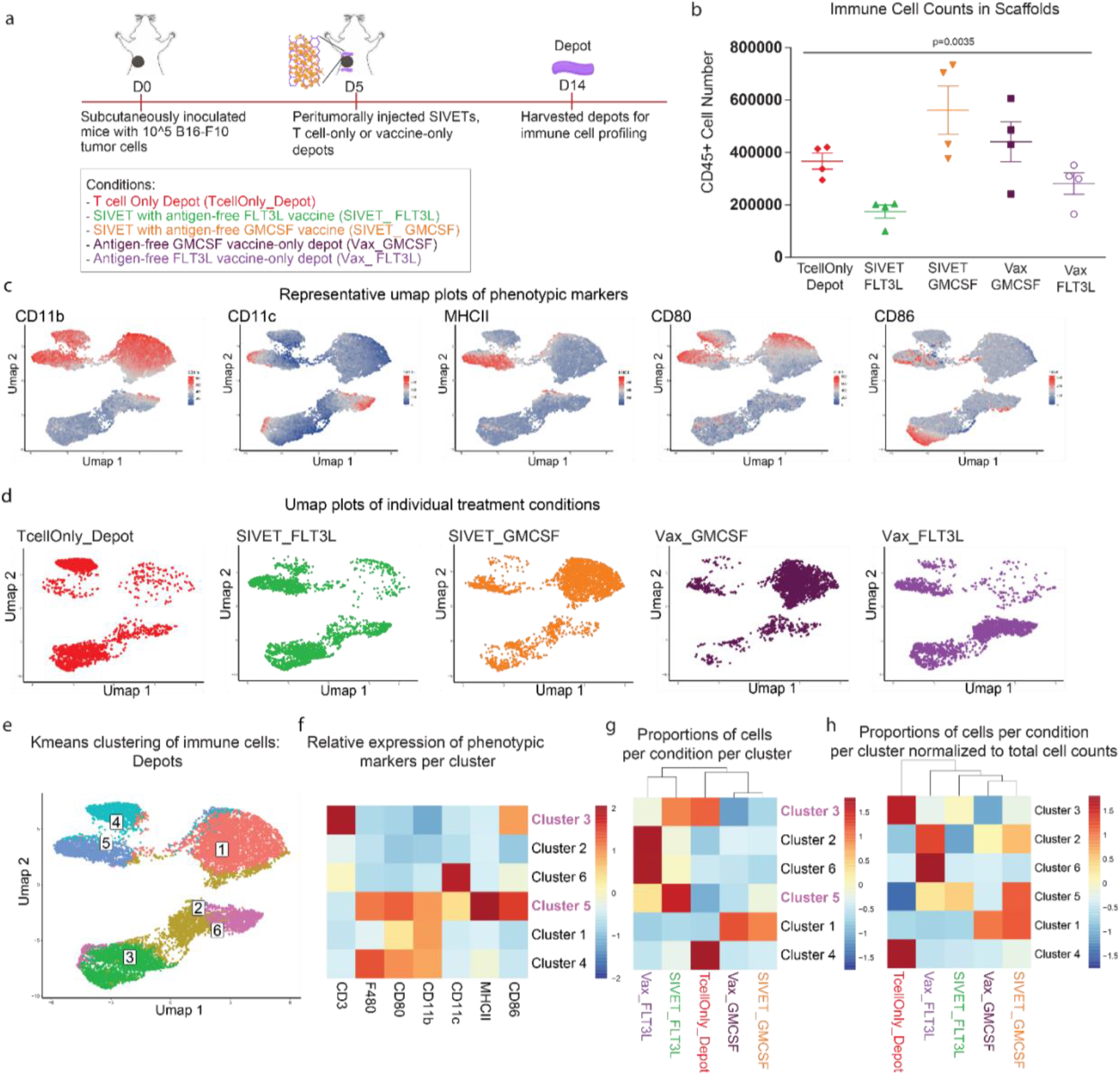
SIVETs enhance the relative levels of activated antigen-presenting cells in depots. a. Schematic of experiment. b. Total counts of immune cells that infiltrated depots per condition. p-values determined by two-tailed one-way ANOVA with Geisser-Greenhouse correction. Data are mean ± s.e.m from n=4 mice per condition. c. Umap plots showing expression of the indicated markers. d. Umap plots of individual treatment conditions showing distinct localization of cells based on treatment group. e. Umap plot overlaid with Kmeans clusters of immune cells in depots. f. Heatmap plot showing the average expression of the indicated immune cell markers in each cluster after K-means analysis. g. Heatmap plot showing the proportion of cells in each condition represented in each cluster. Some clusters of interest are highlighted. Data are pooled cells from n=3 mice per condition. h. Heatmap plot showing the proportion of cells in each condition represented in each cluster, normalized to the total cell counts in (b). Data are pooled cells from n=3 mice per condition.

**Supplementary Fig. 5.**
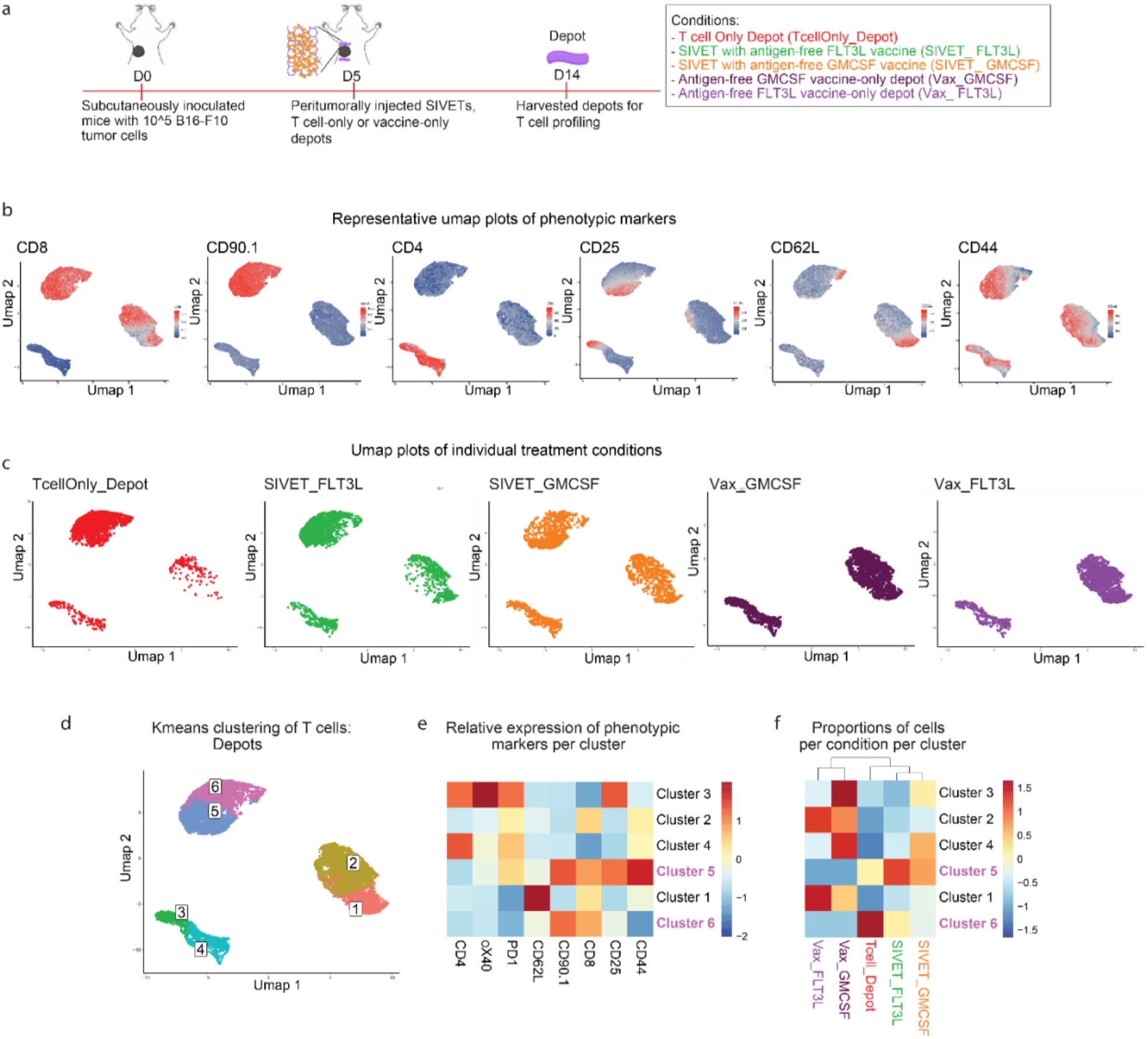
SIVETs prolong activation of adoptively transferred T cells in depots. a. Schematic of experiment. b. Umap plots showing expression of the indicated markers. c. Umap plots of individual treatment conditions showing distinct localization of cells based on treatment group. d. Umap plot overlaid with Kmeans clusters of T cells in depots. e. Heatmap plot showing the average expression of the indicated T cell markers in each cluster after K-means analysis. f. Heatmap plot showing the proportion of cells in each condition represented in each cluster. Some clusters of interest are highlighted. Data are pooled cells from n=3 mice per condition.

**Supplementary Fig. 6.**
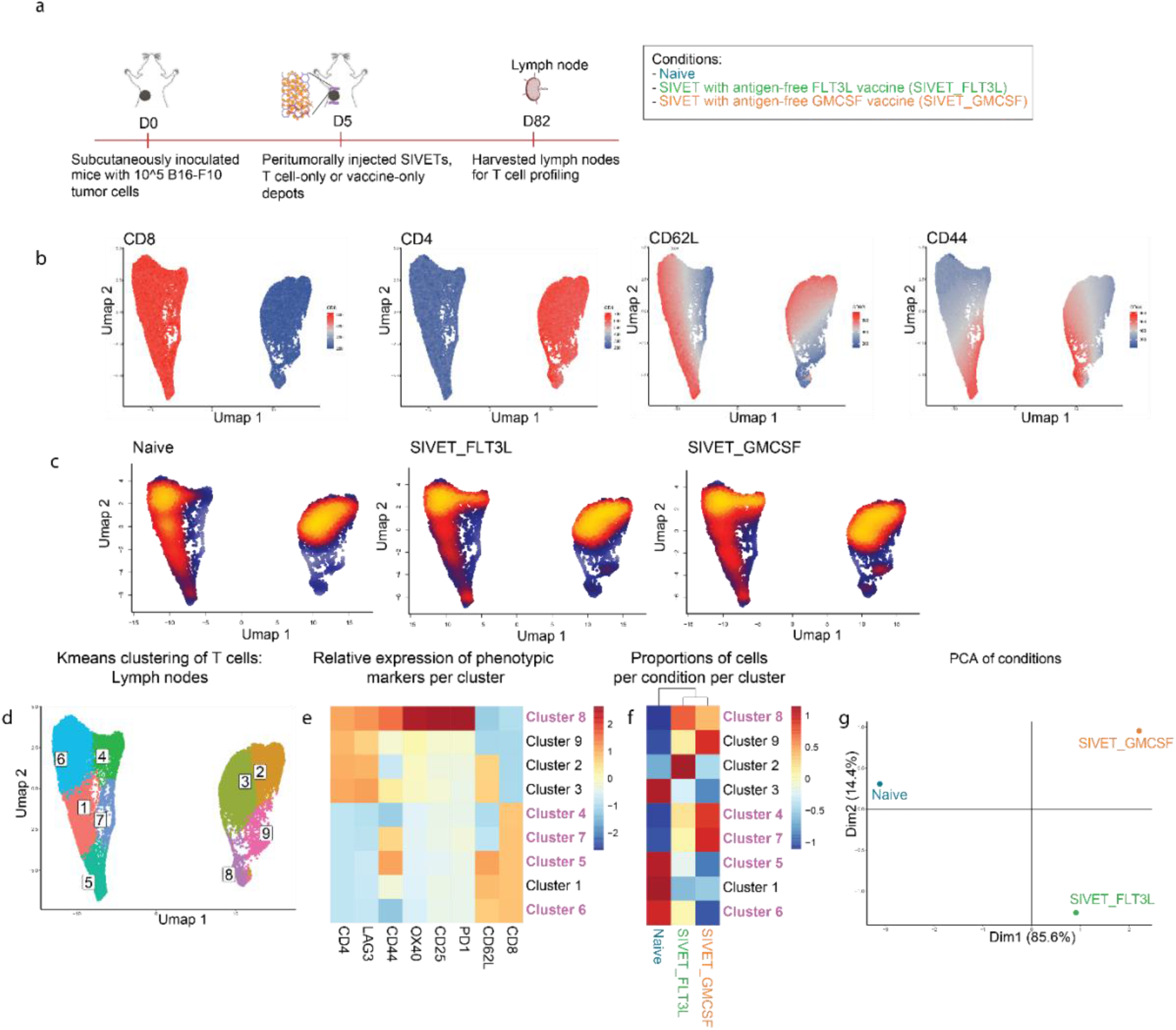
Characterization of long-term T cell profiles in lymph nodes after SIVET treatment. a. Schematic of experiment. b. Umap plots showing expression of the indicated T cell markers. c. 2D density plots showing umap of individual treatment conditions. Denser (hot) regions indicate more cells. d. Umap plot overlaid with Kmeans clusters of T cells in lymph nodes. e. Heatmap plot showing the average expression of the indicated T cell markers in each cluster after K-means analysis. f. Heatmap plot showing the proportion of T cells in each condition represented in each cluster. Some clusters of interest are highlighted. g. PCA plot showing relative similarities between the different treatment conditions. Data are pooled cells from n=3 mice per condition (SIVET_FLT3L and SIVET_GMCSF) and n=2 mice for naïve controls.

**Supplementary Fig. 7.**
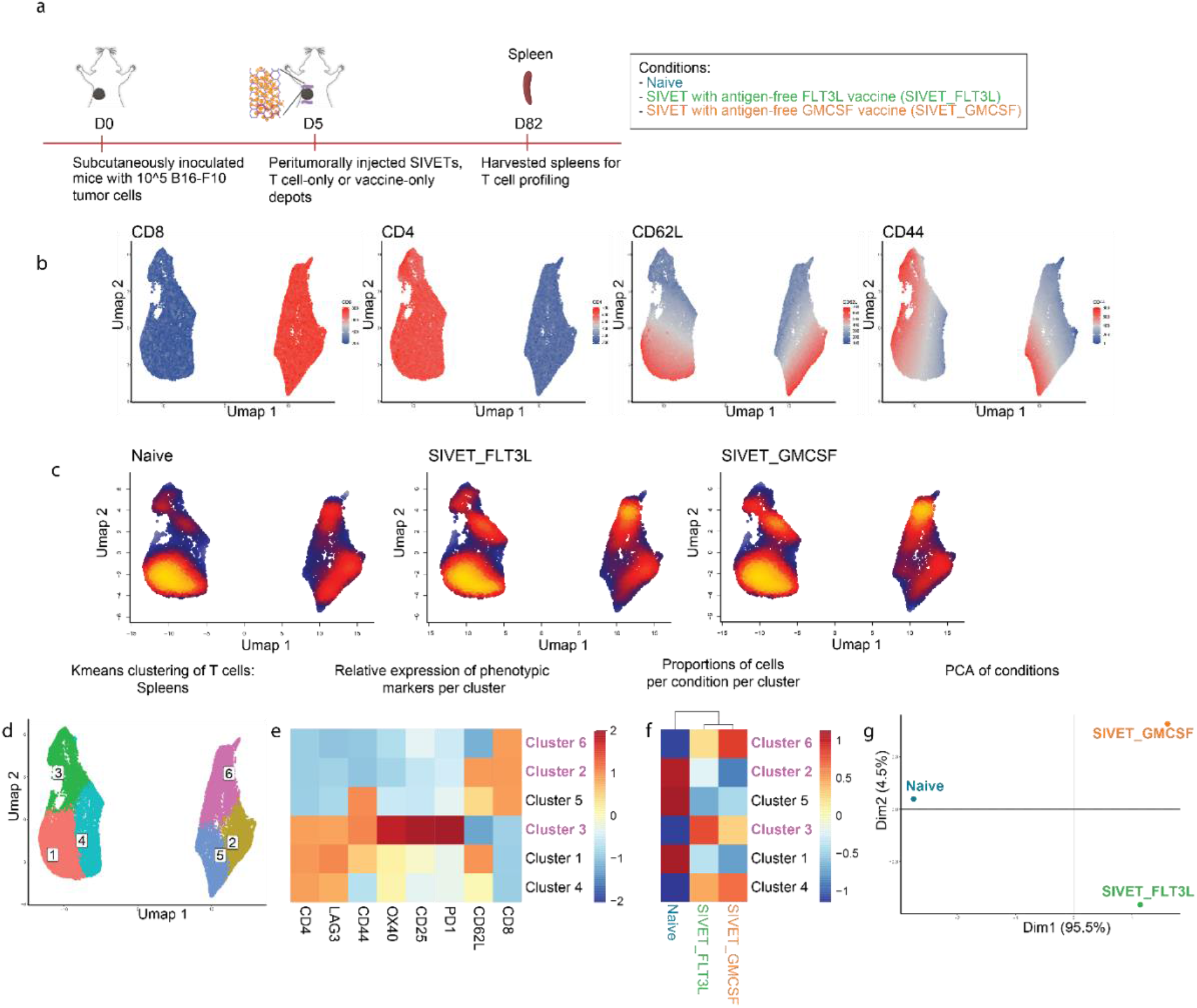
Characterization of long-term T cell profiles in spleens after SIVET treatment. a. Schematic of experiment. b. Umap plots showing expression of the indicated T cell markers. c. 2D density plots showing umap of individual treatment conditions. Denser (hot) regions indicate more cells. d. Umap plot overlaid with Kmeans clusters of T cells in spleens. e. Heatmap plot showing the average expression of the indicated T cell markers in each cluster after K-means analysis. f. Heatmap plot showing the proportion of T cells in each condition represented in each cluster. Some clusters of interest are highlighted. g. PCA plot showing relative similarities between the different treatment conditions. Data are pooled cells from n=3 mice per condition (SIVET_FLT3L and SIVET_GMCSF) and n=2 mice for naïve controls.

**Supplementary Fig. 8.**
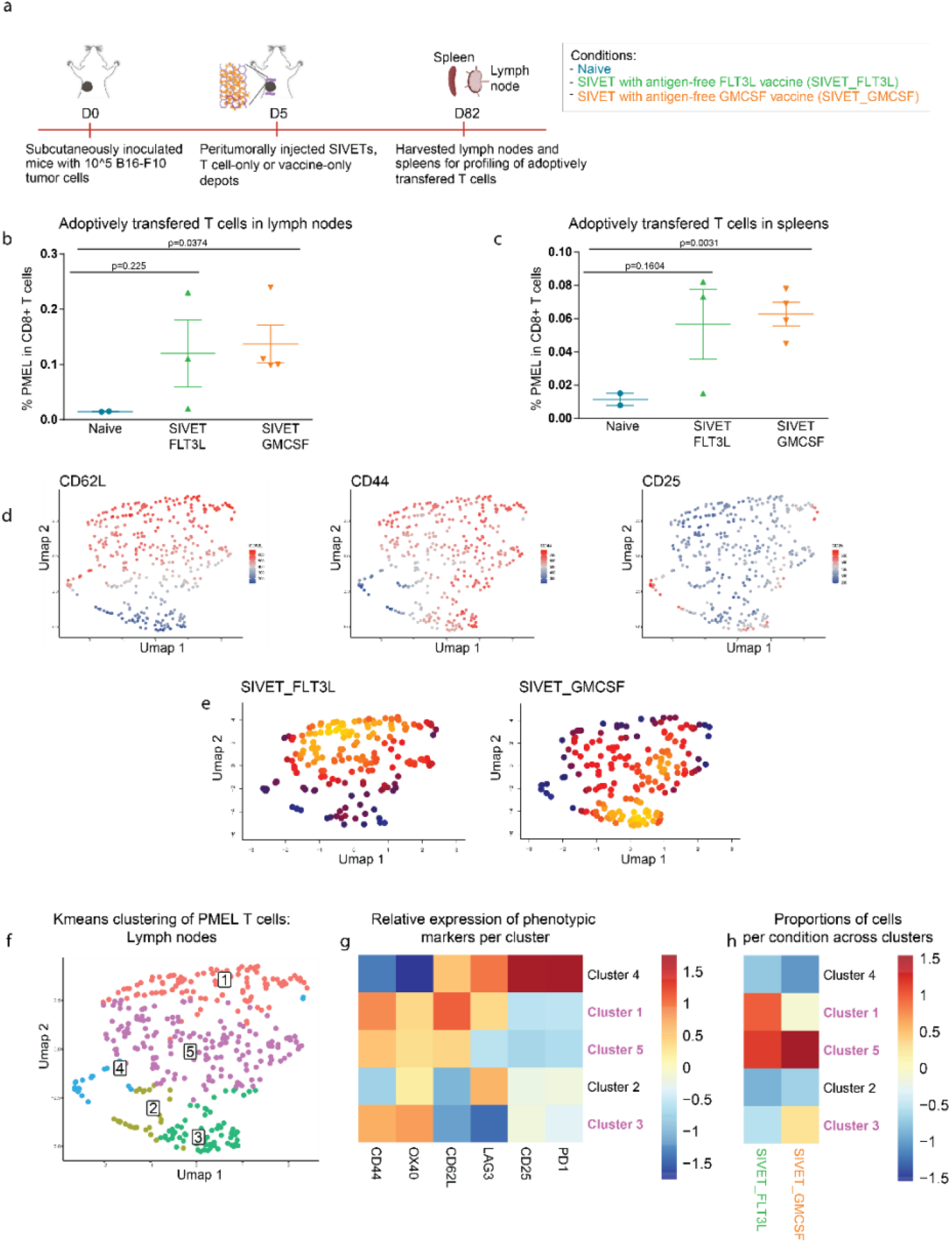
Characterization of long-term profiles of adoptively transferred T cells after SIVET treatment. a. Schematic of experiment. b-c. Proportions of adoptively transferred T cells in CD8+ populations in lymph nodes (b) and spleens (c) of long-term surviving mice. p-values determined by two-tailed unpaired t test with Welch’s correction. Data are mean ± s.e.m from n=3 mice per condition (SIVET_FLT3L and SIVET_GMCSF) and n=2 for naïve. d. Umap plots of adoptively transferred T cells from lymph nodes showing expression of the indicated T cell markers. e. 2D density plots showing umap of individual treatment conditions. Denser (hot) regions indicate more cells. f. Umap plot overlaid with Kmeans clusters of T cells. g. Heatmap plot showing the average expression of the indicated T cell markers in each cluster after K-means analysis. h. Heatmap plot showing the distribution of cells in each condition across the various clusters. Some clusters of interest are highlighted. Data are pooled cells from n=3 mice per condition.

**Supplementary Fig. 9.**
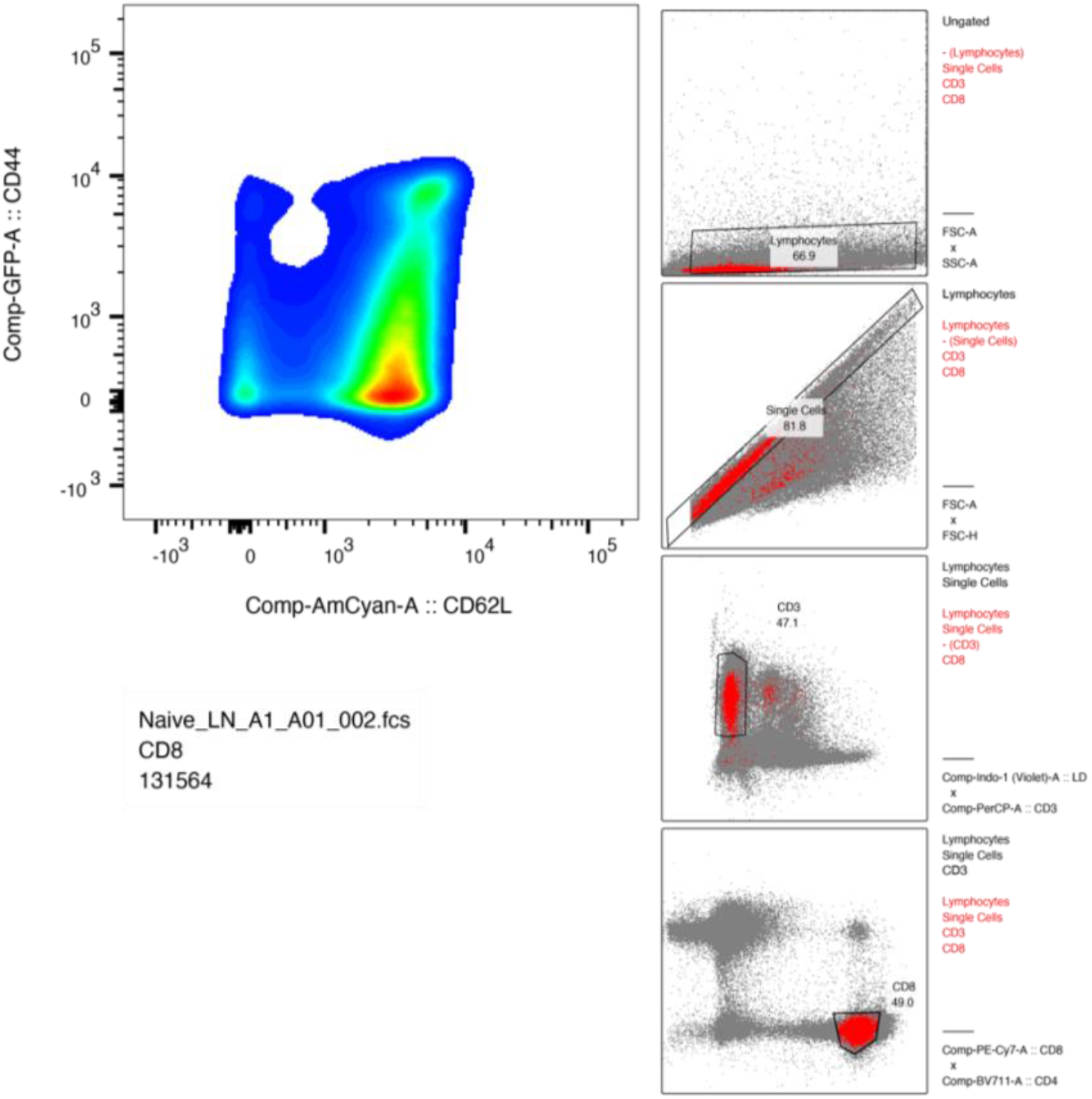
Flow cytometry gating strategy. Representative flow cytometry gating strategy showing steps from ungated population to CD62L and CD44 CD8^+^ populations.

## Acknowledgements

We acknowledge funding from the National Institutes of Health (U54 CA244726, U01 CA214369), the National Science Foundation (MRSEC DMR-1420570), and the Food and Drug Administration (R01 FD006589). The contents are those of the author(s) and do not necessarily represent the official views of, nor an endorsement, by the funding agencies. SEM was performed at the Center for Nanoscale Systems (CNS) at Harvard. Some of the schematics were made using Biorender. We would like to thank the Wucherpfennig lab for their kind gift of B16-cOVA cell line. We would like to acknowledge Dr. Alexander Tatara and Nikko Jeffreys for their scientific inputs. We also thank the staff at the Wyss Institute for Biologically Inspired Engineering at Harvard University for providing the support needed to perform the required experiments, including T. Ferrante, M. Perez, G. Cuneo, E. Zigon, M. Rousseau and M. Carr.

## Correspondence

Correspondence to David J. Mooney

